# ZCCHC4 Promotes Translation of Replication-dependent Histone mRNAs by Recruiting Cytoplasmic eIF3 complex

**DOI:** 10.1101/2025.06.21.660898

**Authors:** Ruiqi Wang, Xiaoyan Shi, Yangyi Zhang, Yuci Wang, Yanlan Cao, Rui Feng, Chen Chen, Yingchun Zhang, Hao Chen, Honghui Ma

## Abstract

Chromatin instability is a major driving force in tumorigenesis, so the correct chromatin structure is essential for maintaining genome stability during the various stages of the cell cycle and DNA template processes. However, there is limited research on how histones are regulated and whether insufficient histone supply leads to chromatin instability, thereby affecting the process of tumor generation.

Our study demonstrates that ZCCHC4 exhibits dual subcellular localization in both the nucleus and cytoplasm. Notably, in the cytoplasm, ZCCHC4 facilitates the translation of replication-dependent (RD) histone mRNAs, potentially through direct interaction with components of the translation initiation machinery. Depletion of ZCCHC4 results in a global reduction in protein translation, particularly affecting the translation of replication-dependent histones. This translational impairment prolongs the S phase of the cell cycle, leading to insufficient histone supply, chromatin relaxation, and genomic instability. Moreover, ZCCHC4 knockout markedly inhibits cell proliferation, colony-forming ability, and the growth of xenograft tumors.

In conclusion, this study demonstrates that ZCCHC4 plays a pivotal role in regulating global protein translation by interacting with the eIF3 complex, which is essential for efficient mRNA translation in cancer cells and for sustaining the supply of replication-dependent histones during the S phase.

## INTRODUCTION

The transformation of normal cells into cancer cells typically involves several steps, usually including the activation of oncogenes and the inactivation of tumor suppressor genes and pro-apoptotic genes ^1^.In many cases, the inactivation of genes crucial for cancer development occurs through epigenetic silencing, typically involving hypermethylation of CpG-rich promoter regions. Hypermethylation of CpG islands in gene promoter regions maintains gene silencing at the transcriptional level, thereby suppressing the expression of downstream genes^2,3^. This leads to cellular dysfunction and increases the risk of cancer.

The RAS family of oncogenes is associated with the immortalization of many cell lines, and mutations in RAS genes are present in approximately 30% of human tumors. In 2007, *Nature* magazine reported that Michael R. Green and colleagues, through RNAi library screening, discovered 9 epigenetic modification-related genes closely associated with anchorage-independent growth and tumor growth suppression in K-RAS transformed NIH3T3 cells^4^. ZCCHC4 is one of these genes. Hepatitis B virus (HBV) infection poses a serious threat to our health. The immune evasion mechanisms of HBV in both its virology and immune response remain not fully understood. In 2017, *Cell* magazine reported a study by Professor Cao Xuetao’s team, where they conducted an RNAi screen of 771 epigenetic regulatory molecules in the HepG2.2.15 cell line infected with HBV^5^. The study identified 37 genes that affect the secretion of HBsAg, including SETD2 and ZCCHC4. Further validation with multiple different shRNAs confirmed that SETD2, ZCCHC4, and others inhibit the secretion of HBsAg. Previous studies demonstrated that ZCCHC4 catalyzes m⁶A methylation at position A4220 of 28S rRNA^6^, the basic research on ZCCHC4 is crucial for understanding tumor formation and viral immune mechanisms. However, the specific mechanism by which ZCCHC4 affects tumor formation in the K-RAS pathway is currently unknown.

Chromatin instability is a major driving force in tumorigenesis, so the correct chromatin structure is essential for maintaining genome stability during the various stages of the cell cycle and DNA template processes^7,8^. However, current research on histones largely focuses on histone post-translational modifications (PTMs), histone variants, nuclear and nucleolar interactions, and ATP-dependent chromatin remodeling. However, there is limited research on how histones are regulated and whether insufficient histone supply leads to chromatin instability, thereby affecting the process of tumor generation. Classical histone genes are expressed in a replication-dependent (RD) manner. Replication-dependent histones are unique in terms of structure, content, and expression.

The mRNA of replication-dependent histones contains a stem-loop structure at its 3 end, consisting of a six-base stem and a four-nucleotide loop, instead of a poly(A) tail^9,10^. The stem-loop sequence is capable of binding to the stem-loop binding protein (SLBP). The stem-loop sequence is followed by the histone downstream element (HDE), which base pairs with U7 small nuclear RNA (snRNA). This interaction plays a crucial role in the stability, degradation, and translation efficiency of mRNA, particularly during different stages of the cell cycle. The content of histones in the cell is proportional to the DNA content, and their synthesis is tightly regulated during cell division. Histone synthesis peaks during the S phase, ensuring that newly synthesized DNA can be packaged into chromatin along with histones. Histone plays a decisive role in the structure of chromosomes by helping DNA form nucleosomes and participating in higher-order chromatin structures. They regulate gene expression by altering the accessibility of genes.

Histone translation is particularly active during the S phase, and cell cycle regulatory proteins ensure the sufficient supply of histones during DNA replication by regulating their synthesis^11^. The S phase is the peak period for histone expression. Like most mRNAs, replication-dependent histone mRNA is exported from the nucleus via the antigen peptide transporter, allowing for efficient translation^12–14^. The translation of polyadenylated mRNA requires the 3’poly(A) tail to be brought near the 5’cap. This is mediated by protein-protein interactions between poly(A)-binding protein and eukaryotic translation initiation factor 4G (EIF4G), which also interacts with the cap-binding protein^15^. Replication-dependent histone mRNA circularization has developed a unique mechanism^16^, where the stem-loop binding protein (SLBP) first binds to the stem-loop structure of the histone mRNA^9,17^. Subsequently, SLBP interacts with another protein, SlIP1, which also interacts with the translation initiation factor EIF4G. Through these interactions, SLBP and SlIP1 bring the 3’ end of histone mRNA closer to the 5’cap structure, thereby forming a circular structure^18^. This circularization mechanism is similar to the translation mechanism of polyadenylated mRNA, both of which promote efficient translation through protein-protein interactions. Therefore, SLBP plays a crucial role in the circularization of histone mRNA^19,20^. It is not only responsible for binding to the specific structure of the mRNA but also facilitates mRNA translation through interactions with other proteins. At the end of the S phase, a short U-tail is added to the 3’ end of histone mRNA^21^. The U-tail is bound by the LSM1-7 ring complex, which helps recruit the recapping complex and exonuclease complexes, leading to the degradation of the mRNA in both the 5’ to 3’ and 3’ to 5’ directions^22,23^. Additionally, cyclin A (Cyc A) and cyclin-dependent kinase 1 (CDK1) phosphorylate SLBP, triggering its degradation at the end of the S phase, thereby preventing further histone mRNA synthesis^24,25^.

In contrast, a small group of histone variants are replication-independent (RI), primarily H3.3 and H2AZ. These variants are present in all multicellular organisms, their mRNA is expressed throughout the cell cycle, and they have polyA tails^10^. For example, mammalian cells express a large amount of Histone H3 variants, including H3.1 and H3.3, which play important regulatory roles in somatic cells. The typical H3.1 is primarily expressed during the S phase and is considered to be replication-coupled, while H3.3 is expressed throughout the cell cycle and is replication-independent. Histone genes are generally highly conserved, but in some cases, certain histone variants may lead to distinct gene expression patterns.

## RESULTS

### ZCCHC4 affects global translation by recruiting eIF3 complex

It has been demonstrated that ZCCHC4 catalyzes m⁶A methylation at position A4220 of 28S rRNA^6,26^, a process that primarily occurs in the nucleus^27^. Intriguingly, we found that ZCCHC4 exhibits nucleo-cytoplasmic dual localization^6^ in HeLa cells, prompting us to further investigate its potential involvement in biological processes occurring in the cytoplasm. Firstly, we validated the nucleo-cytoplasmic dual localization of ZCCHC4 through cell fractionation analysis in both HeLa and PLC/PRF/5 cells (Figure. 1a), which indicated that ZCCHC4 is almost evenly distributed in cytoplasm and nucleus. To explore its potential function in cytoplasm, we employed the tandem affinity purification coupled with mass spectrometry (TAP-MS) in ZCCHC4-overexpressing cells to identify its interacting proteins (Figure. 1b). Our mass spectrometry analysis confirmed interactions between ZCCHC4 and both eIF3 subunits and its associated factors, most notably PDCD4^28^ (Figure. 1c). Through co-immunoprecipitation (Co-IP) assays, we further validated the specific interaction between ZCCHC4 and eIF3H, a core subunit of the eIF3 complex. Importantly, the binding between these two proteins remained stable even after RNase A treatment. This result demonstrates that the ZCCHC4-EIF3H interaction likely represents a direct protein-protein interaction, rather than an RNA-mediated indirect binding event (Figure 1d). Previous structural study showed that ZCCHC4 contains multiple domains: an N-terminal GRF zinc finger domain (residues 24–100), followed by a C2H2 zinc finger domain (residues 100–163), a methyltransferase (MTase) domain (residues 163–378), and a C-terminal CCHC zinc finger domain (residues 378–464)^29^, which are folded into an integrated functional module (Extended Data Figure. 1a). To identify the domains responsible for ZCCHC4 binding to the eIF3 complex, we generated a series of domain-truncated mutants in 293T cells and performed Co-IP assays. The results demonstrated that only the full-length ZCCHC4 could stably bind to eIF3H and PDCD4, whereas individual truncated domains failed to efficiently interact with the eIF3 complex (Extended Data Figure. 1b). This suggests that the interaction between ZCCHC4 and eIF3 likely depends on the integrity of its overall structure rather than the independent participation of individual domains.

**Fig. 1:**
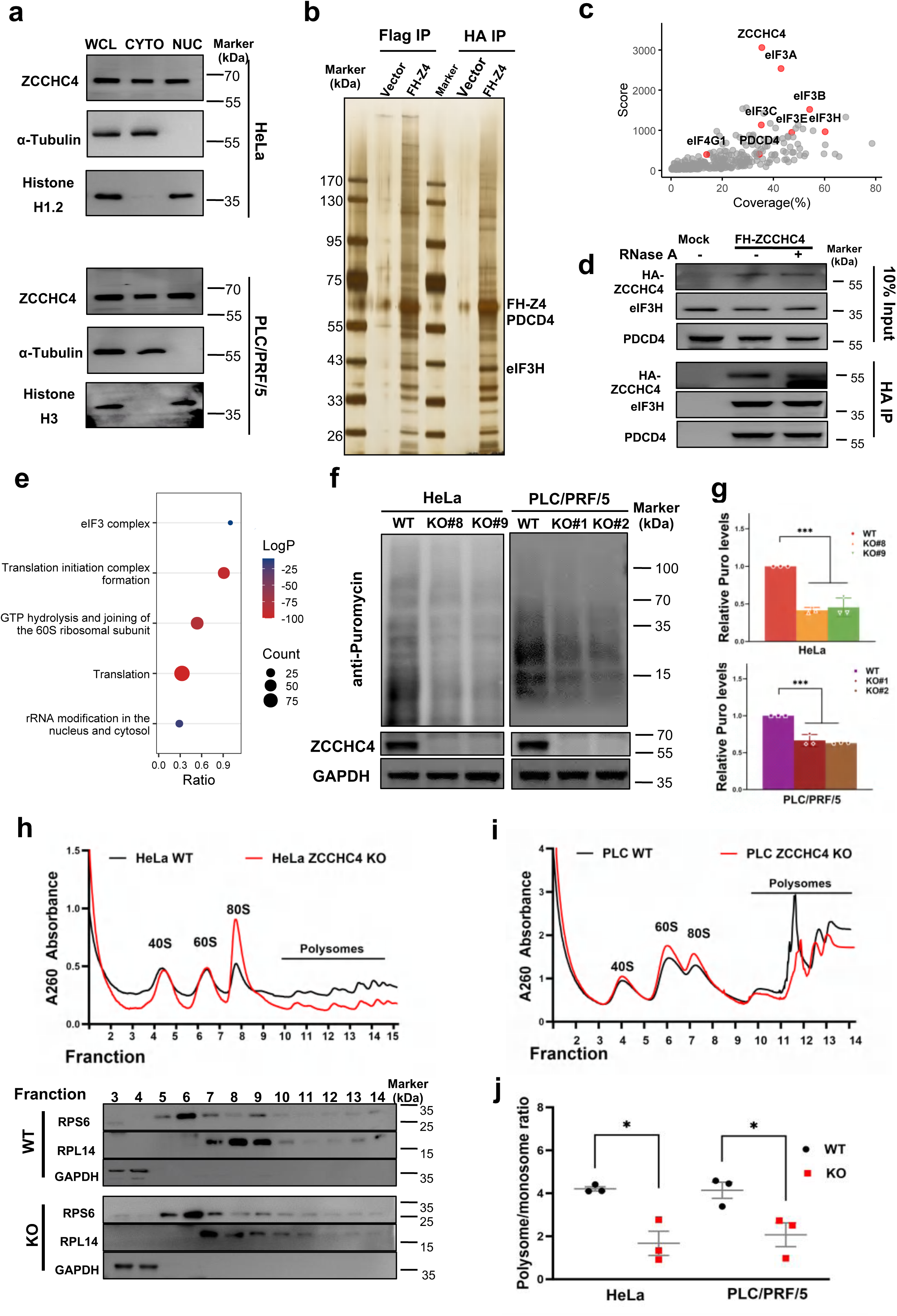
ZCCHC4 regulates global mRNA translation through interaction with the eIF3 complex. **a,** Western blot analysis of ZCCHC4 in the subcellular fractions of HeLa and PLC/PRF/5 cells. Tubulin was used as a cytoplasmic marker, Histone H1.2 and Histone H3 as the nuclear markers. **b,** Identification of ZCCHC4 interacting partners. ZCCHC4 complex was purified from ZCCHC4-FH stable cells using immunoaffinity purification. Eluted proteins were separated by SDS-PAGE and analyzed by silver staining. **c,** ZCCHC4 associates with the multiple subunits of the eIF3 complex (highlighted by red dots) in mass spectrometry analysis. **d,** Western blot confirmed that eIF3H and PDCD4 are the binding proteins of ZCCHC4. **e,** GO analysis of ZCCHC4-associated proteins identified by mass spectrometry. **f, g,** Global protein synthesis in ZCCHC4 knockout (KO) and control cells was measured by puromycin assay in Hela cells (**f,** left) and PLC/PRF/5 (**f,** right) cells. Quantification of protein expression levels is shown in (**g**). **h-j,** Polysome profiling were showed the change of global translation after ZCCHC4 knockout in Hela cells (**h,** up) and PLC/PRF/5 cells (**i**). Western blot confirms fractionation integrity in Hela cells using 40S (RPS6) and 60S (RPL14) ribosomal markers, with GAPDH as a ribosome-free loading control (**h,** down). The experiment was repeated three times, and the ratio of poly- to mono- (AUC) was statistically analyzed (**j**). **Note: g**, **j**, One-way ANOVA with Tukey’s test. \**P*<0.05, \*\*\**P*<0.001. Data were presented as mean ± SD (n=3).

Considering that the eIF3 complex plays a central role in translation initiation regulation through its interaction with the 40S ribosomal subunit^30,31^, we systematically analyzed the binding properties of ribosomal components via sucrose gradient centrifugation (Extended Data Figure. 1c). The experiment separated the small ribosomal subunit (40S), large ribosomal subunit (60S), intact ribosomes (80S), and polysome fractions, then examined the distribution patterns of ZCCHC4, EIF3A, EIF3B, and EIF3H. The results demonstrate that ZCCHC4 not only colocalizes with the eIF3 complex in certain fractions but is also widely present across all ribosomal fractions, including the polysome fraction (Extended Data Figure. 1d). These findings suggest that ZCCHC4 may regulate protein translation by interacting with the eIF3 complex. Consistently, the marked enrichment of ZCCHC4-interacting proteins in the cytoplasmic translation pathway (Figure. 1e), suggests that its cytoplasmic localization may directly contributes to mRNA translation regulation. Therefore, we performed polysome analysis and puromycin experiments to investigate the changes in translation in HeLa and PLC/PRF/5 cells after ZCCHC4 knockout. Stable ZCCHC4 knockout cell lines were generated by CRISPR-based genetic recombination technology (Extended Data Figure. 1e, f), and the depletion of ZCCHC4 was validated by western blot and RT-qPCR analysis (Extended Data Figure. 1g, h). Puromycin assay demonstrated that ZCCHC4 knockout significantly reduced puromycin incorporation into nascent polypeptides (Figure. 1f, g), while polysome profiling analysis revealed a marked decrease in polysome fractions upon ZCCHC4 depletion (Figure. 1h-j). These results indicate that ZCCHC4 plays an important regulatory role in global protein synthesis in HeLa and PLC/PRF/5 cells. Taken together, the above data demonstrated that ZCCHC4 is likely a critical regulator of mRNA translation in cancer cells, where it modulates global protein synthesis via direct engagement with the eIF3 complex.

### ZCCHC4 specifically regulates the translation of replication-dependent histone mRNAs

To define the mRNA translation categories modulated by ZCCHC4, we first re-analyzed our PAR-CLIP-seq (photoactivatable ribonucleoside cross-linking and immunoprecipitation coupled with high-throughput sequencing) to identify direct mRNA targets of ZCCHC4^6^. In two biological replicates, a total of 340 mRNAs that interacted with ZCCHC4 were identified, among which 28 histone mRNAs were found to be targeted by ZCCHC4 (Figure. 2a). Analysis of these 28 histone mRNAs revealed that ZCCHC4 primarily binds to the CDS regions of target mRNAs through the CAA motif (Figure. 2b). To validate the ZCCHC4 target mRNAs identified by PAR-CLIP-seq, we selected four histone transcripts-H2B, H2AC, H1B, and H4A-for further validation by RIP-qPCR analysis. Consistently, we found significant enrichment of these mRNAs in anti-ZCCHC4 immunoprecipitates from HeLa cells, confirming the possible targeting by ZCCHC4 (Figure. 2c).

**Fig. 2:**
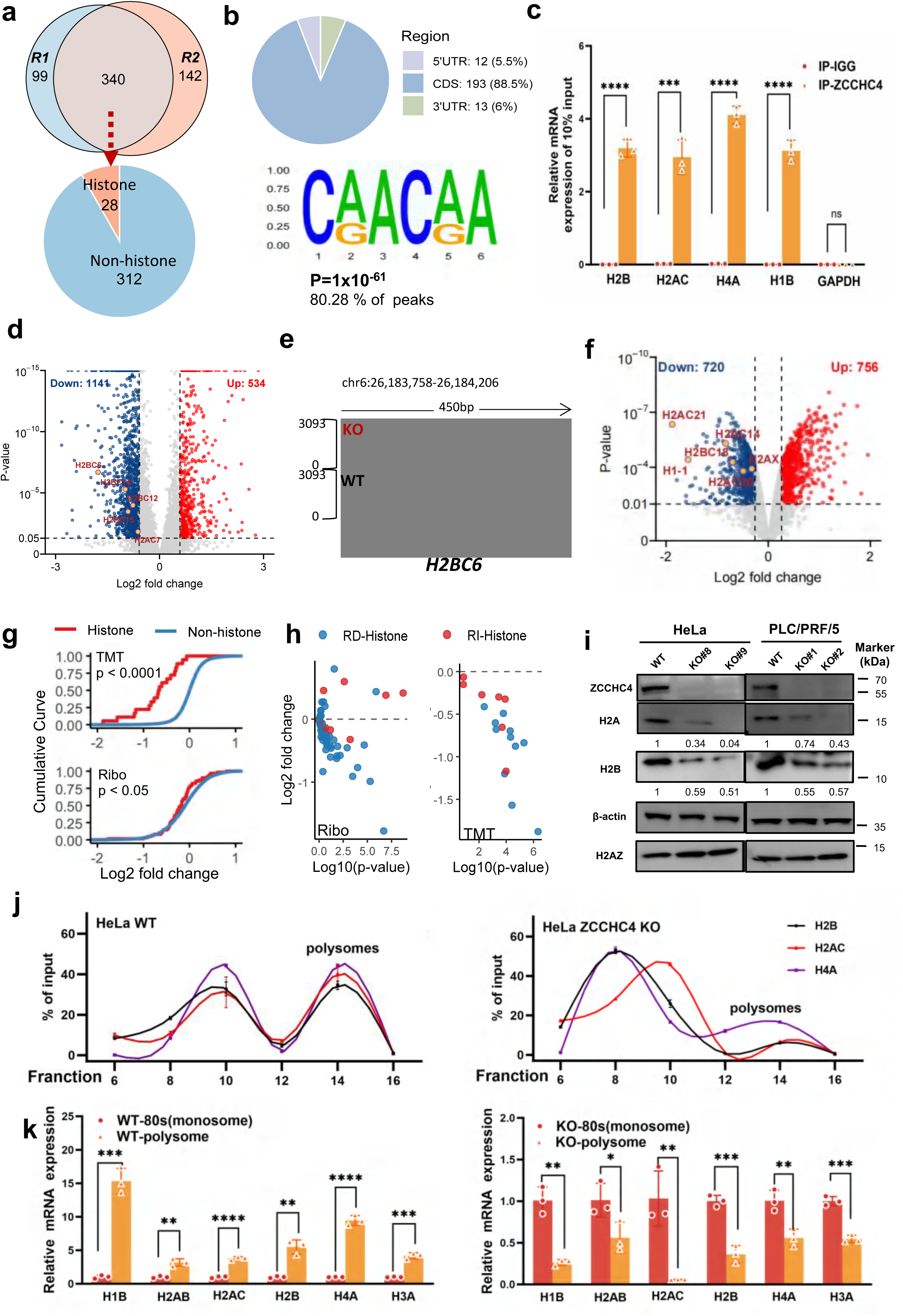
ZCCHC4 promotes the translation of histone mRNA targets. **a,** Venn diagram showed the overlap of ZCCHC4 PAR-CLIP-seq target genes between two biological replicates. **b,** Distribution plots (up) and sequence motif enrichment analysis (down) of ZCCHC4-bound histone mRNAs. ZCCHC4-binding motifs were identified by HOMER findMotifsGenome.pl from the PAR-CLIP-seq peaks of two biological replicates. The motif length was restricted to 4-8 nucleotides. The *P*-value was calculated from the random background sequences with a ZOOPS score. **c,** RIP/qPCR analyses showed the binding of ZCCHC4 and potential targets in wild-type HeLa cells. **d,** Volcano plot depicting down-regulated (blue) and up-regulated (red) genes identified by Ribo-seq in ZCCHC4 KO cells compared with WT cells. **e,** Integrated Genome Viewer (IGV) showing the mapped reads of the binding peaks identified by ribosome-bound fragments identified by Ribo-seq of H2BC6 after ZCCHC4 knockout. The histograms represent the sum of mapped reads along the genome. The mapped reads of Ribo-seq were normalized by the sequencing depth. **f,** Volcano plot depicting down-regulated (blue) and up-regulated (red) genes identified by TMT in ZCCHC4 KO cells compared with WT cells. **g,** Cumulative distribution of translation efficiency with histone mRNA (red) and non-histone (blue) mRNA by TMT (up) and Ribo-seq (down). **h,** Volcano plot depicting replication-dependent (blue, RD-Histone) and replication-independent (red, RI-Histone) histone identified by Ribo-seq (left) and TMT (right) in ZCCHC4 KO cells compared with WT cells. **i,** Western blot analysis to validate the expression of RD-histone (H2A and H2B) and RI-histone (H2AZ) in ZCCHC4 knockout and control cells. β-actin was used as internal control. **j,** The broken line chart showed the relative distribution of the mRNA level of H2B, H2AC and H4A in the different fractions isolated by the sucrose gradient. The same volume (50 µL) of the fractions was loaded to perform RT-qPCR. **k,** RT-qPCR analysis of polysome-or monosome (80S)-bound histone mRNA in WT (left) ZCCHC4 knockout (right) HeLa cells. **Note: c**, **k** Two-tailed unpaired Student’s t tests. Data were presented as mean ± SD (n=3). \**P*<0.05, \***\****P*<0.01, \*\***\****P*<0.001, \*\*\*\**P*<0.0001. ns: not significant.

To gain deeper insights into the role of ZCCHC4 in the regulation of protein synthesis, we compared changes in the translation efficiency (TE) of ZCCHC4-targeted transcripts between ZCCHC4 knockdown cells and control cells by ribosome footprinting (Ribo-seq) (Extended Data Figure. 2a-c). We identified 1675 differentially translated genes with high confidence (Ribo-seq transcripts per million > 1, RNA-seq transcripts per million > 5, fold change >= 1.5, and p-value < 0.05). Among the translationally altered genes, there are 1141 genes with decreased translation efficiency and 534 genes with increased translation efficiency in the ZCCHC4 KO cells compared with the WT cells (Figure. 2d). Histone mRNAs were substantially enriched among downregulated genes (Figure. 2e). To further characterize the protein-level changes influenced by ZCCHC4, we performed TMT (tandem mass tag)-based quantitative proteomic analysis towards WT and ZCCHC4 KO cells (Extended Data Figure. 2d). A total of 6703 proteins were identified, among which 720 were significantly downregulated and 756 were upregulated (Figure. 2f). Of note, the protein levels of H2A and H2B were also significantly reduced in ZCCHC4 KO cells among the detected histones (Figure. 2f, Extended Data Figure. 2e). To determine whether ZCCHC4 influences histone translation or protein levels, we compared the TE and expression of histone-containing versus non-histone transcripts. Notably, the translation efficiency of histone mRNAs and histone protein level were preferentially affected by ZCCHC4 knockout (Figure. 2g). It’s well-documented that the histones can be divided into two categories: replication-dependent histones (RD) and replication-independent histones (RI) based on different gene expression regulatory mechanisms. Interestingly, we found that RD histones were more significantly affected (Figure. 2h).

To validate the Ribo-seq and TMT proteomics data, we performed western blot and RT-qPCR analyses on selected histones. Consistent with multi-omics results, the protein levels of RD-histones (H2A and H2B) were significantly reduced in ZCCHC4 knockout cells, whereas the RI-histone H2A.Z showed no significant change (Figure. 2i). Notably, these changes occurred without corresponding alterations in histone mRNA levels (Extended Data Figure. 2f), suggesting that ZCCHC4 primarily regulates gene expression of histone at the translational rather than transcriptional level. Furthermore, we performed polysome profiling of ribosome-bound fractions. Strikingly, histone mRNA levels in polysome fractions were significantly higher in control cells compared to ZCCHC4 KO cells, while the opposite trend was observed in 80S monosome fractions (higher in KO cells) (Figure. 2j). This clear shift from polysome to monosome association in ZCCHC4-KO cells demonstrates ZCCHC4 depletion impaired ribosomal loading and reduced translational efficiency. Importantly, these results were further confirmed by RT-qPCR analysis of key ribosome profiling fractions, providing independent validation of the translational defect (Figure. 2k). Taken together, these data support that ZCCHC4 specifically regulates the translation of RD histone mRNAs without affecting their transcription.

### Deficiency of histone in ZCCHC4-KO cell leads to loosened chromatin and interrupted cell cycle

The synthesis of histones is tightly coordinated with DNA replication to ensure proper chromatin assembly and genome stability. Given that ZCCHC4 regulates the translation of histone mRNAs, its deficiency may cause insufficient histone supply, thereby disrupting nucleosome assembly and affecting chromatin structure. To investigate whether ZCCHC4 knockout indeed influences chromatin accessibility, we conducted a comparative analysis of ATAC-seq data between WT and ZCCHC4 KO cells (Extended Data Figure. 3a), thoroughly examining the impact of ZCCHC4 depletion on chromatin accessibility and its association with histone homeostasis. We found that ZCCHC4 knockout significantly altered chromatin architecture, inducing a global increase in accessibility, where 288 gene loci exhibited enhanced openness while 36 regions became more restricted (Figure. 3a). Importantly, transcription start sites (TSS) displayed stronger chromatin-protection signals in WT cells compared to ZCCHC4-KO counterparts (Figure. 3b, Extended Data Figure. 3b), suggesting that ZCCHC4 depletion induces chromatin destabilization at regulatory regions. The precise regulation of histone mRNA synthesis and turnover is critical for proper S-phase progression in the cell cycle^32^. Canonical histone supply directly influences DNA synthesis rates, and insufficient histone levels extend S-phase duration, ensuring accurate replication and facilitating the G2/M transition^32^. This delay likely arises from impaired replication fork progression due to reduced histone availability^33,34^. Given that ZCCHC4 mediates histone translational regulation and the tight coupling between histone dynamics and cell cycle control, we hypothesized that ZCCHC4 may mechanistically regulate cell cycle progression. Consistent with this notion, our TMT-based quantitative proteomics analysis revealed significant enrichment of downregulated proteins in both cell cycle regulation and histone assembly pathways (Extended Data Figure. 3c).

**Fig. 3:**
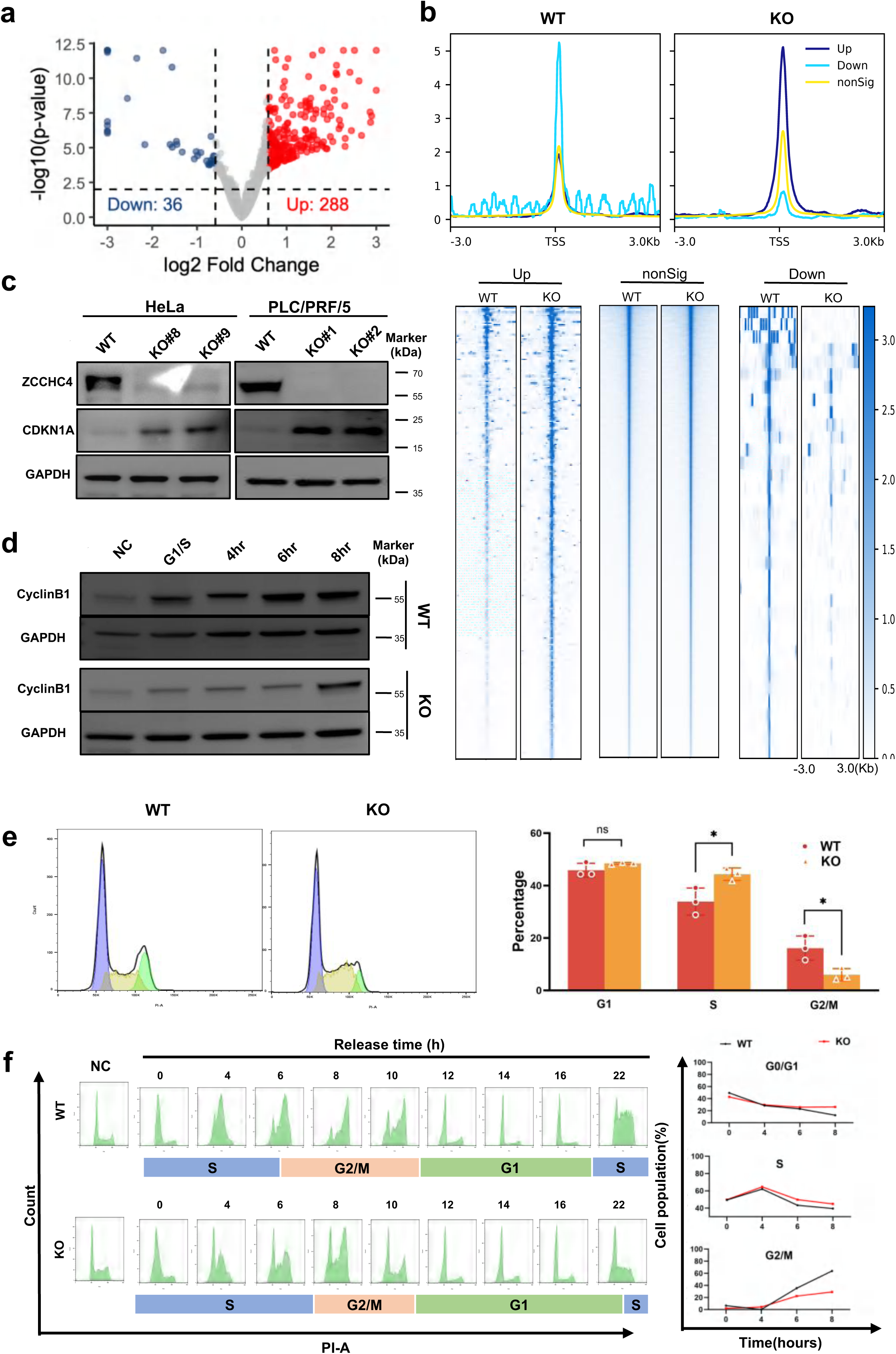
ZCCHC4 depletion induces cell cycle arrest in S phase. **a,** Volcano plot depicting down-regulated (blue) and up-regulated (red) genes identified by ATAC-seq in ZCCHC4 KO cells compared with WT cells. **b,** Accessible chromatin profiling of HeLa WT and ZCCHC4 KO cells by ATAC-Seq. Peak distributions are presented as a profile plot (upper panel) and heatmap (low panel) over ± 3 kb centered on the TSSs. **c,** Western blot of CDKN1A in HeLa WT and ZCCHC4 KO cells. GAPDH was used as a loading control. **d,** WT and ZCCHC4 knockout cells were subjected to a double thymidine block to arrest cells at the G1/S checkpoint (G1/S). Cells were released from the block for either 4 or 8h and immunoblotted with the cyclinB1 antibodies to determine the expression of cyclin proteins. Async, asynchronous cells. GAPDH was used as a loading control. Blots shown are representative of three independent experiments. **e,** Representative images (left) and quantification (right) of cell cycle progression of ZCCHC4 KO versus WT HeLa cells. **f,** Propidium iodide staining detected by flow cytometry showing the distribution of each cell cycle stage (G1, S, and G2/M phase) after ZCCHC4 knockout at 0, 4, 6, 8, 10, 12, 14, 16 and 22h after cell synchronization and unsynchronized cells. The flow cytometry profiles (left) and quantitative statistical curves (right) are presented. **Note: e,** Two-tailed unpaired Student’s t tests. Data were presented as mean ± SD (n=3). \**P*<0.05, ns: not significant.

To test this hypothesis, we analyzed cell cycle alterations in ZCCHC4 knockout versus wild-type cells by Western blot. The results showed significantly upregulated CDKN1A expression upon ZCCHC4 depletion, suggesting cell cycle dysregulation (Figure. 3c). Subsequently, cell cycle profiling by flow cytometry (quantified phase proportions based on the Watson pragmatic model^35^) demonstrated significant S-phase prolongation concomitant with G2/M-phase contraction in ZCCHC4-knockout cells (Figure. 3e). In contrast, ZCCHC4 depletion does not affect cell apoptosis (Extended Data Figure. 3d). To precisely delineate ZCCHC4’s role in cell cycle regulation, we performed G1/S-phase synchronization via double-thymidine blockade in isogenic HeLa WT and ZCCHC4 KO cell lines, followed by cell cycle progression analysis. The thymidine blockade was validated to arrest cells at the G1/S boundary (G1/S arrest), and upon release from the block, cells progressed through the S phase into G2. Following release from the arrested state, we also examined the expression of cell cycle-related proteins using western blot analysis. In control cells, the level of cyclin B began to increase at 6 hours post-release, indicating entry into mitosis. However, this increase was delayed until 8 hours in ZCCHC4-depleted cells, suggesting a delayed S phase progression in the absence of ZCCHC4 (Figure. 3d). Following release from the thymidine-induced arrest, we assessed cell cycle status 22 hours post-DNA staining using the Watson pragmatic model. The results showed that ZCCHC4-depleted cells exhibited delayed cell cycle progression, remaining in the S phase while control cells had already progressed to the G2/M phase (Figure. 3f). In summary, these data indicate that ZCCHC4 is essential for maintaining histone supply during the S phase.

### ZCCHC4 is necessary for tumor growth

Given the critical role of cell cycle regulation in tumorigenesis, we then asked whether ZCCHC4 will play a critical role in tumorigenesis. ZCCHC4 deletion exerted a strong suppression of cell growth in both PLC/PRF/5 and Hela cells (Figure. 4a, b). Then, monoclonal formation assay and wound-healing assay were carried out, and we observed that depletion of TARBP1 decreased both colony-forming abilities and migratory abilities of cancer (Figure. 4c-e). To further investigate the role of ZCCHC4 in cell proliferation, we performed an EdU incorporation assay to evaluate DNA synthesis activity. The results showed that ZCCHC4 KO cells exhibited a marked reduction in the proportion of EdU-positive cells, indicating impaired S phase progression and decreased DNA replication capacity. These findings support a critical role for ZCCHC4 in regulating cell cycle progression and maintaining proliferative activity (Figure. 4f).

**Fig. 4:**
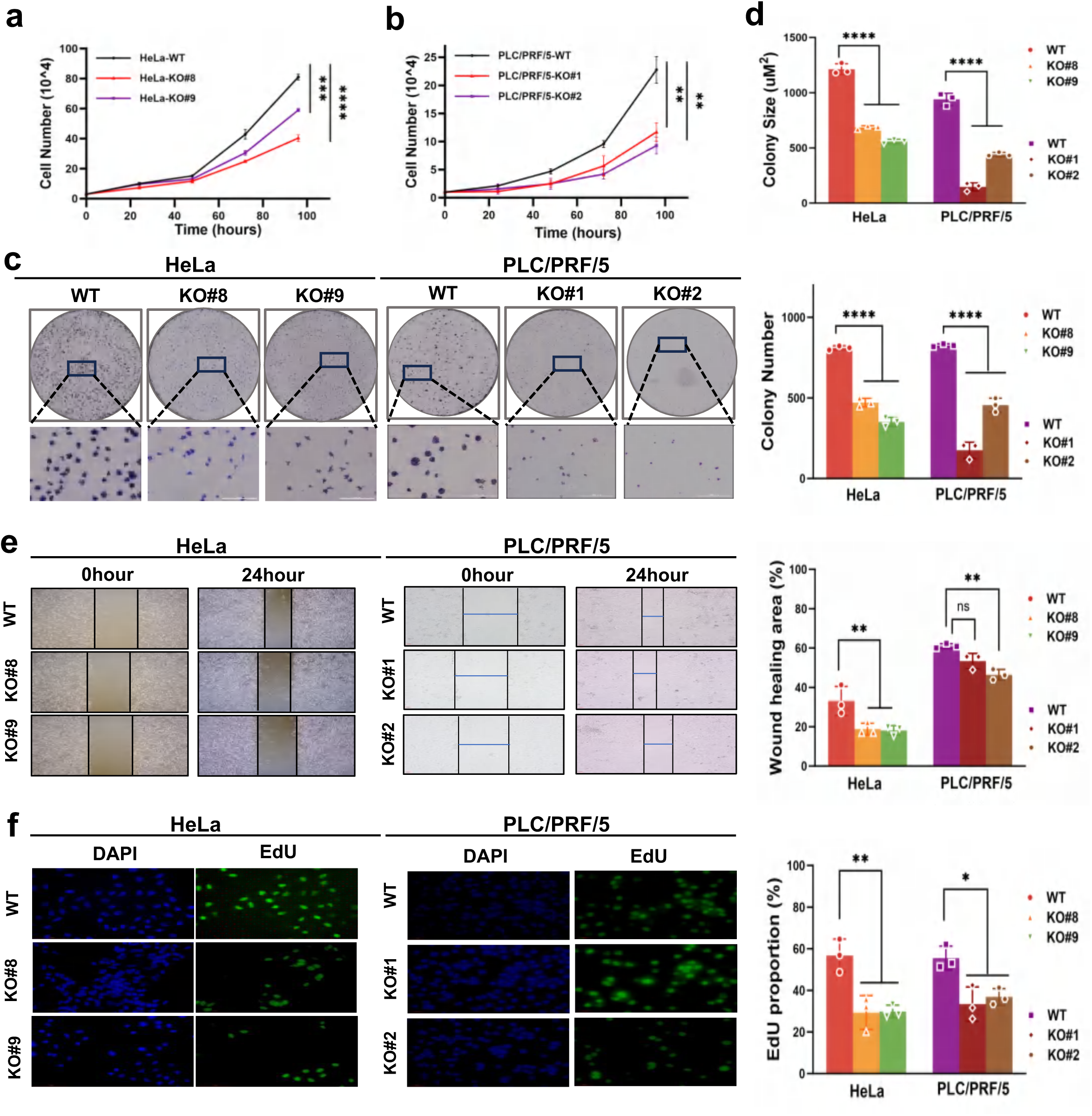
ZCCHC4 promotes cancer cell growth, survival and invasion. **a, b,** Cell proliferation is dramatically repressed upon ZCCHC4 knockout. In HeLa cell line (**a**), 3 × 10^4^ cells were seeded in one well of a 24-well plate on day 0 and counted each day until day 3 with cell counters. In PLC/PRF/5 cell line (**b**), 1 × 10^4^ cells were seeded in one well of a 24-well plate on day 0 and counted on day one until day 3 with cell counters. **c,** Colony-formation assay of WT and ZCCHC4 KO in HeLa and PLC/PRF/5 cells. 1000 cells were seeded in 6-cm tissue culture dishes, representative pictures were taken (left), and the colony numbers and sizes (**d**) were measured on day 9 (right). **e,** Wound-healing assay of WT and ZCCHC4 KO in HeLa and PLC/PRF/5 cells. Representative pictures were taken (left), and the scratch area after healing was measured at 24 hours (right). **f,** EdU proliferation assay showing cell proliferation of ZCCHC4 cells after ZCCHC4 knockout in HeLa and PLC/PRF/5 cells. Representative images (left) and quantitative statistical analysis (right) are shown. Green indicated the EdU positive cells. Note: **a, b** Two-way ANOVA. **c, d, e, f** One-way ANOVA with Tukey’s test. **a-f** Data were represented as mean ± SD (n = 3). \**P* < 0.05, \*\**P* < 0.01, \*\*\**P* < 0.001, \*\*\*\**P* < 0.0001, ns: not significant.

To conclude, our findings demonstrate that ZCCHC4 is essential for tumor growth. To test whether these tumorigenic effects are specifically mediated by ZCCHC4, we performed genetic rescue experiments by reintroducing ZCCHC4 into ZCCHC4-depleted cells (KO+RE) (Extended Data Figure. 5a) and observed a partial rescue of cellular proliferation (Figure. 5a), colony formation (Figure. 5b, c), and xenograft tumor growth compared to KO cells (Figure. 5d-f). Consistently, immunohistochemical staining for PCNA and Ki67 revealed that ZCCHC4 re-expression partially restored proliferative marker levels (Figure. 5g), while BCL2 staining remained largely unchanged across all groups (Extended Data Figure. 5b), indicating minimal impact on apoptosis. Taken together, our study proves that ZCCHC4 specifically recruits the eIF3 complex to regulate the translation of RD-histones, thereby mediating cell cycle control in cancer cells and promoting tumorigenesis and progression (Figure. 5h).

**Fig. 5:**
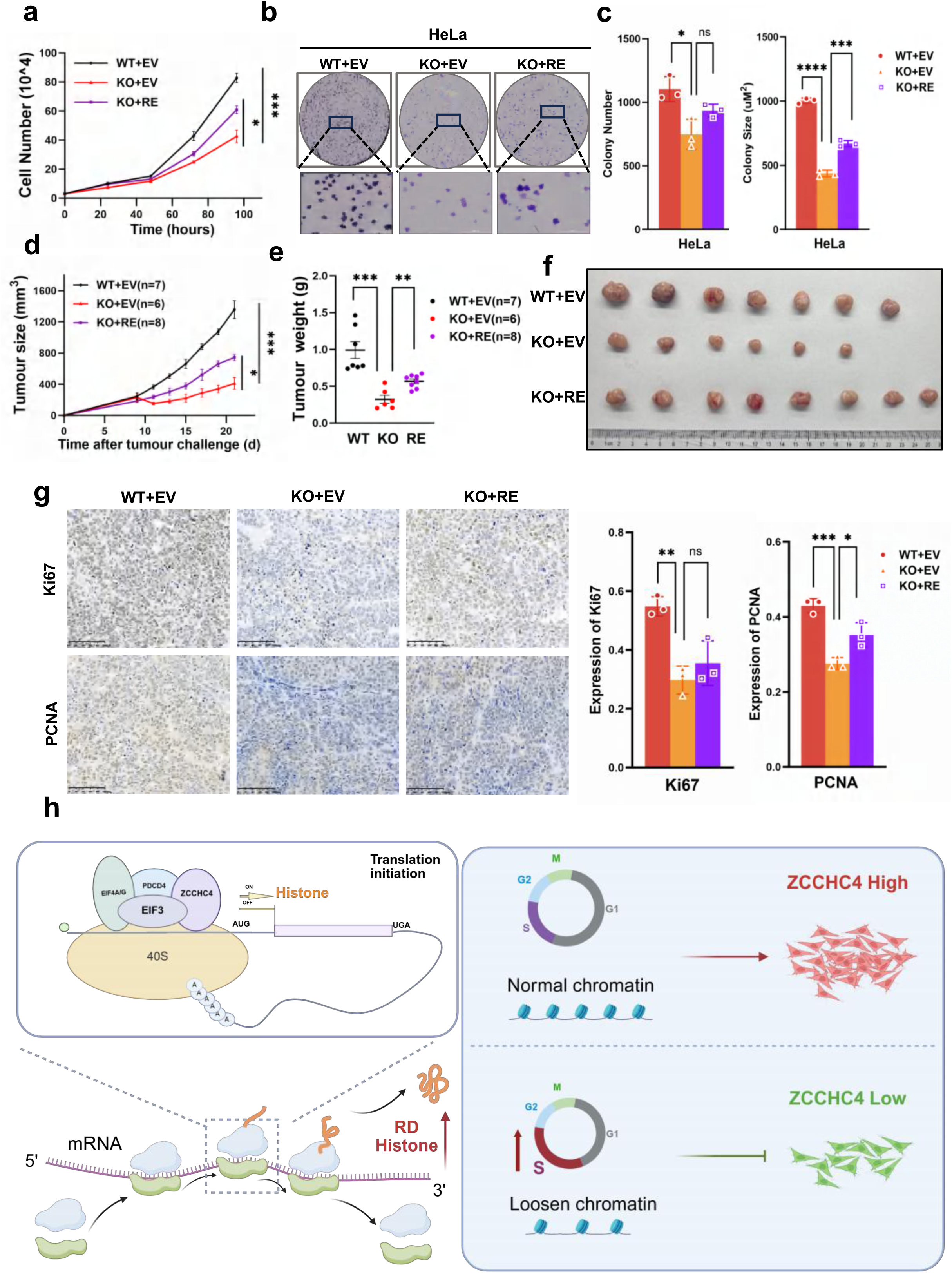
ZCCHC4 is necessary for tumor growth. **a-f,** Overexpression of ZCCHC4 partially rescues cell proliferation(**a**), colony formation (**b, c**), and xenograft tumor growth (**d-f**) of ZCCHC4 KO cells. 3 × 10^4^ cells were seeded in one well of a 24-well plate on day 0 and quantified every 24 h until day 4 with cell counters (**a**). 1000 cells were seeded in a 6-well plate and cultured for six days (**b**). The tumor growth (**d**) and final tumor weights (**e**) were compared, and a picture of the final tumors was taken (**f**). **g,** Immunohistochemistry expression of Ki67 and PCNA from WT, ZCCHC4 KO and KO rescued tumor. **h,** Working model for the critical role of ZCCHC4 in tumor. ZCCHC4 promotes translation of replication-dependent histone mRNAs by recruiting the cytoplasmic eIF3 complex, in turn enhancing cell cycle progression and driving tumorigenesis. Note: **a, d** Two-way ANOVA. **c, e, g** Two-tailed unpaired Student’s t tests. **a, c, g,** Data were represented as mean ± SD(n=3). **d, e,** Data were presented as mean ± SEM. \**P* < 0.05, \*\**P* < 0.01, \*\*\**P* < 0.001, \*\*\*\**P* < 0.0001, ns: not significant.

## DISCUSSION

In the present study, we identified ZCCHC4 as a novel regulator of cell cycle progression through its role in modulating histone mRNA translation. ZCCHC4 is an m⁶A RNA methyltransferase previously known to catalyze the methylation of 28S rRNA, thereby maintaining ribosome function. Consistent with its dual localization in both the nucleus and cytoplasm, our data suggests that ZCCHC4 may also exert non-catalytic functions by interacting with components of the eIF3 complex. Notably, knockout of ZCCHC4 significantly impairs global protein synthesis, with a pronounced reduction in the translation of replication-dependent histone mRNAs, leading to S phase prolongation, chromatin assembly defects, and attenuated cell proliferation. These findings establish ZCCHC4 as a key player in coordinating protein synthesis with cell cycle demands. Given the essential role of histone production in sustaining rapid DNA replication and chromatin organization, ZCCHC4-mediated translational control may contribute to tumor cell proliferation. Overall, our study reveals the oncogenic potential of ZCCHC4 through its regulation of histone biosynthesis and cell cycle dynamics, providing new mechanistic insight into its role in cancer progression.

### Histone translation regulation

Histone synthesis is a tightly regulated process that must be precisely coordinated with DNA replication to ensure genome integrity^32^. During the S phase, cells require a sharp increase in histone production to package newly synthesized DNA into chromatin^9,10^. Unlike most proteins regulated transcriptionally, replication-dependent (RD) histone genes are primarily controlled at the post-transcribing and translational levels^36^. These transcripts lack poly(A) tails and contain a stem-loop structure in the 3′ UTR, allowing phase-specific control through RNA-binding proteins such as SLBP^10^. By contrast, replication-independent (RI) histones are expressed throughout the cell cycle and are involved in chromatin remodeling, transcriptional regulation, and DNA repair^37^.

In our study, we found that knockout of ZCCHC4 selectively impairs the translation of RD histone mRNAs, while having minimal effects on RI histones. This finding suggests that ZCCHC4 plays a critical role in coupling histone synthesis with cell cycle progression. Our data suggests it may influence mRNA translation by interacting with the eIF3 translation initiation complex. Given that RD histone mRNAs require precise translational control and possess unique structural elements, ZCCHC4 might function as a scaffold protein that facilitates their selective recruitment to the ribosome.

Clinically, elevated RNA polymerase II (RNAPII) occupancy at histone gene loci has been associated with higher histone transcription, accelerated chromatin assembly, and poor prognosis in multiple cancers^38^. This observation supports the notion that aberrant histone expression facilitates tumor cell proliferation. We speculate that ZCCHC4 may contribute to this oncogenic process by promoting the translation of histone mRNAs or other chromatin assembly-related transcripts, thereby sustaining rapid cell division in tumor settings.

Taken together, our findings identify ZCCHC4 as a key regulator of replication-dependent histone translation and cell cycle progression.

### Non-catalytic functions

RNA methyltransferases are key epigenetic regulators that primarily catalyze methyl modifications on various types of RNA molecules (e.g., m⁶A, m⁶Am, m⁵C). These modifications influence RNA stability, splicing, nuclear export, degradation, and translation, thereby playing essential roles in post-transcriptional regulation, cell fate determination, stress response, and tumorigenesis^5,39–41^. For instance, m⁶A modification can affect ribosome recruitment and initiation complex assembly, directly modulating the translation efficiency of target mRNAs and reshaping the cellular proteome. The underlying mechanism is likely mediated through its regulation of rRNA methylation, which modulates ribosome function. This is consistent with the previous findings by Ma *et al*., who reported that ZCCHC4 catalyzes m⁶A modification at position A4220 of 28S rRNA, thereby influencing the translation process^6^.

Increasing evidence, however, suggests that certain RNA methyltransferases also possess non-catalytic, structural functions. METTL3, for example, has been shown to enhance translation of target oncogenic mRNAs by directly interacting with the eIF3 complex in the cytoplasm, independent of its catalytic activity^42^. Similarly, METTL16 has been reported to act as a scaffold protein involved in mRNA splicing and nuclear export^43^. Although the study by Ma *et al*. suggested that ZCCHC4-mediated m⁶A modification of 28S rRNA primarily affects ribosome structure and 60S subunit assembly^6^, Pinto *et al*. reported that knockout of ZCCHC4 does not substantially impair ribosome maturation or assembly^26^. These differing observations suggest that additional regulatory mechanisms may be involved.

Our study shows that ZCCHC4, in addition to its established role as an rRNA m⁶A methyltransferase localized in the nucleolus, is also partially localized in the cytoplasm, where it physically interacts with the eIF3 translation initiation complex. Given the central role of eIF3 in selective mRNA translation initiation, we speculate that ZCCHC4 may exert non-catalytic regulatory functions like METTL3^42^, acting as a structural component of translational complexes and selectively promoting the translation of specific transcripts such as replication-dependent histone mRNAs. This hypothesis provides a potential explanation for how ZCCHC4 contributes to translational regulation and cell cycle progression independently of its methyltransferase activity.

### ZCCHC4 axis in tumorigenesis

In recent years, the oncogenic potential of ZCCHC4 has garnered increasing attention across various cancer types. Studies have reported elevated expression of ZCCHC4 in malignancies such as small cell lung cancer, colorectal cancer, and esophageal cancer, where its overexpression correlates strongly with poor prognosis and chemoresistance^44–46^. Our experimental data further demonstrate that knockout of ZCCHC4 significantly suppresses tumor cell proliferation, colony formation capacity, and xenograft tumor growth, underscoring its critical role in promoting tumorigenesis.

This tumor-promoting effect may result from ZCCHC4-mediated enhancement of the translation of specific oncogenic and metabolic transcripts, which facilitates sustained proliferation and invasiveness of cancer cells under high metabolic stress^47,48^. The roles of ZCCHC4 in other physiological processes, including stem cell differentiation and immune responses, merit deeper exploration^49–51^.

Moreover, ZCCHC4 regulates the translation efficiency of specific mRNAs, such as key S-phase regulatory proteins, through interaction with eIF3, ensuring adequate protein synthesis for rapid cell proliferation. This mechanism may also participate in cancer cells’ responses to environmental stress and chemotherapeutic agents. Previous studies have implicated ZCCHC4 in cisplatin resistance via the ROS/c-MYC axis^46^, and our results further support its role in post-transcriptional translational regulation. Thus, ZCCHC4 represents a promising therapeutic target for overcoming chemoresistance in cancer treatment.

In summary, this study identifies ZCCHC4 as a novel translational regulator of replication-dependent histone mRNAs, playing a critical role in S-phase progression and tumor cell proliferation. Mechanistically, ZCCHC4 possesses catalytic activity for m⁶A modification on rRNA and may also enhance the translation of histone mRNAs by interacting with the cytoplasmic eIF3 complex. By coordinating ribosome function with the selective translation of cell cycle regulators, ZCCHC4 ensures timely histone supply during DNA replication. Moreover, ZCCHC4 is frequently overexpressed in various cancers and is significantly associated with poor prognosis and chemoresistance. This study reveals a previously unrecognized mechanism of translational control mediated by ZCCHC4 and highlights its potential as a promising therapeutic target in cancer treatment.

## METHODS

### Cell culture

HeLa cells, PLC/PRF/5 cells and HEK 293T cells were purchased from Wuhan Pricella Biotechnology. All cells were authenticated by STR profiling and testing for mycoplasma contamination was performed with a MycAway one-step mycoplasma detection kit (Yeasen, 40612ES25). All cells were cultured in Dulbecco’s Modified Eagle Medium (DMEM) supplemented with 10% fetal bovine serum (FBS, BI) and 1% penicillin-streptomycin (15-140-122; Thermo Fisher Scientific, Waltham, MA, USA). The cells were maintained in a humidified incubator at 37°C with 5% CO₂. Cells were passaged every 2-3 days using 0.25% trypsin-EDTA to detach them from the culture flask. For experiments, cells were seeded at an appropriate density and allowed to adhere overnight before treatment or further analysis.

### Plasmid construction

ZCCHC4 KO cells were generated using a CRISPR–Cas9 approach^52^. To generate *ZCCHC4* knockout HeLa cells, two 20-nt sgRNAs targeting the first/fourth exons of *ZCCHC4* were individually cloned into the pL-CRISPR.EFS.GFP plasmid, while for *ZCCHC4* knockout PLC/PRF/5 cells, two distinct 20-nt sgRNAs targeting the first/second exons were similarly inserted into the same plasmid. This approach ensured efficient CRISPR/Cas9-mediated disruption of *ZCCHC4* in both cell lines. To re-express ZCCHC4 protein into KO cells, ZCCHC4 cDNA was cloned into the pCDH retroviral vector. To overexpress HA and Flag tagged (FH) full-length or FH-truncated ZCCHC4 in HEK 293T cells, the cDNAs encoding full-length or truncated ZCCHC4 (N-terminal GRF zinc finger domain (residues 24-100), MTase domain (residues 163-378), and C2H2-CCHC zinc finger domain (residues 100-163 and 378-464)) was cloned into the pCDH retroviral vector separately.

sgRNAs targeting human *ZCCHC4* locus in HeLa cells

sgRNA#1: GTGCCGGGGAAGCTCGGGAA

sgRNA#2: GGGGCAACATAGTGAGCATC

sgRNAs targeting human *ZCCHC4* locus in PLC/PRF/5 cells

sgRNA#1: CAGGAATGGGTTTGAAGCCG

sgRNA#2: TGTTTGTAAAGGTGACCCAA

### Generation of ZCCHC4 KO cell lines and rescue experiments

To generate *ZCCHC4* knockout cells, 2 μg of sgRNA plasmid was transfected to the corresponding cells using Lipofectamine 3000 (Invitrogen). After 2 days’ culture in DMEM medium, cells were screened according to fluorescence and isolated into monoclons by flow cytometry. After 3 weeks’ culture, colonies were screened out and confirmed by western blot with ZCCHC4 antibody and real-time quantitative PCR analysis.

To generate viruses, the empty vector or FH-ZCCHC4 WT plasmid was co-transfected with pMD2.G and PsPAX2 plasmids into 293T cells, respectively. Hela cells were infected with virus generated by empty vector, whereas KO cells were infected with virus generated by empty vector, FH-ZCCHC4 WT plasmid separately. After 7 days’ selection with 2 μg/ml puromycin, cells were confirmed by western blot with the corresponding antibodies. To overexpress FH-full-length or FH-truncated ZCCHC4 in HEK 293T cells, the full-length or truncated ZCCHC4 plasmid was co-transfected with pMD2.G and PsPAX2 plasmids into 293T cells, respectively. After 7 days’ selection with 2 μg/ml puromycin, cells were confirmed by western blot with the HA antibodies.

### Protein extraction and western blot

HeLa and PLC/PRF/5 Cells were harvested and washed twice with ice-cold phosphate-buffered saline (PBS) to remove any residual media. After centrifugation at 1000 g for 5 minutes, the cell pellets were resuspended in RIPA buffer (R0278; Sigma-Aldrich, St Louis, MO, USA) containing 1% protease inhibitor cocktail (11836153001; Roche, Basel, Switzerland). The cell suspension was incubated on ice for 30 minutes with occasional vortexing to ensure complete cell lysis. Afterward, the lysate was centrifuged at 12000 g for 15 minutes at 4°C to remove debris, and the supernatant was collected for protein quantification. Protein concentration was determined using the BCA kit (23227; Thermo Fisher Scientific). Equal amounts of total protein (30-50µg) were separated by SDS-PAGE (4-20% acrylamide gel) and transferred onto a PVDF membrane (Millipore, IPVH00010). The membrane was blocked with 5% non-fat dry milk in Tris-buffered saline with 0.1% Tween-20 (TBST) for 1 hour at room temperature. The membrane was then incubated with primary antibodies (specific to the protein of interest) overnight at 4°C, followed by incubation with the appropriate HRP-conjugated secondary antibody for 1 hour at room temperature. After washing with TBST, protein bands were visualized using the enhanced chemiluminescence (ECL) detection system (Thermo Fisher Scientific, 32106).

The corresponding antibodies included anti-ZCCHC4 antibody with 1:1000 dilution (Abcam, ab209901), anti-RPS6 antibody with 1:1000 dilution (Proteintech, 66886-1-Ig), anti-RPL14 antibody with 1:1000 dilution (Proteintech, 14991-1-AP), anti-CDKN1A antibody with 1:1000 dilution (Proteintech, 10355-1-AP), anti-puromycin antibody with 1:1000 dilution (ABclonal, A21205), anti-H2B antibody with 1:2000 dilution (PTM-1007), anti-H2A antibody with 1:2000 dilution (ABclonal, A3692), anti-H2AZ antibody with 1:2000 dilution (Proteintech, 16441-1-AP), anti-HistoneH1.2 antibody with 1:1000 dilution (Proteintech, 19649-1-AP), anti HistoneH3 antibody with 1:2000 dilution (ABclonal, A22348), anti-α-Tubulin antibody with 1:3000 dilution (Proteintech, 80762-1-RR), anti-EIF3H antibody with 1:1000 dilution (Proteintech, 11310-1-AP), anti-EIF3A antibody with 1:1000 dilution (Proteintech, 26178-1-AP), anti-PDCD4 antibody with 1:1000 dilution (Proteintech, 12587-1-AP), anti-CyclinB1 antibody with 1:1000 dilution (ABclonal, A19037), anti-GAPDH antibody with 1:5000 dilution (Proteintech, 10494-1-AP), anti-β-actin antibody with 1:3000 dilution (SAB Signalway Antibody, 52901), and anti-Flag antibody with 1:2000 dilution (Sigma, F1804). Full and uncropped western blots are shown in Fig. S6, S7 and S8.

### Cell Proliferation, colony formation, and wound-healing assays

For cell proliferation assays, cells were seeded in individual wells of a 24-well plate on day 0 and counted daily from day 1 to day 3 using an automated cell counter (Thermo Fisher Scientific) to monitor growth rates. For colony formation assays, 1000 cells were seeded in a 6-well plate or 6-cm dish and cultured for six or nine days, respectively. At the endpoint, colonies were fixed with 4% formaldehyde (Beyotime) for 20 minutes and stained with 0.1% crystal violet solution (Solarbio) for 30 minutes. For wound-healing assays, 6×10^4^ cells were seeded in a 6-well plate and cultured in DMEM supplemented with 10% FBS until reaching 90-100% confluency. Cells were then switched to DMEM containing 2% FBS and incubated overnight. Linear scratch wounds were created using a sterile 200 μL pipette tip. Images were taken at 0 h and 24 h post-scratch using a phase-contrast microscope to evaluate cell migration.

### Quantitative PCR and PCR of genomic DNA

Total RNA was extracted using TRIzol reagent (Invitrogen, 15596018). Reverse transcription was performed with 1 mg of total RNA using the HiScript III RT SuperMix for qPCR (Vazyme, R323-01). Quantitative real-time PCR (RT-PCR) was conducted using ChamQ SYBR qPCR Master Mix (Vazyme, Q331-02) on a QuantStudio 3 Real-Time PCR System (Applied Biosystems). Primer sequences are provided in Supplementary Table 1.

### Complex purification

To purify ZCCHC4-associated complex, four 15-cm plates of HeLa cells stably expressing FH-alone (mock) or HeLa cells stably expressing full-length or truncated ZCCHC4–FH were collected and then lysed on ice with 500uL IP buffer (10 mM Tris-HCl pH 7.5, 100 mM KCl, 5 mM MgCl2, 1 mM DTT, 100 μg/ml Cycloheximide, 0.25% NP40) for 10min. Protein extracts were collected via centrifugation at 12000 g for 15 min at 4 °C and then incubated with anti-Flag/HA beads (Invitrogen) overnight for immunoprecipitation. Beads-protein mixture was washed four times with IP buffer before elution, which was performed by competition with flag/HA peptides. Eluted proteins were tested by silver and subsequently subjected to high-performance liquid chromatography-mass spectrometry (HPLC-MS) to analyze ZCCHC4-associated factors.

### SUnSET assay

Cells seeded in 6-well plates were cultured to reach 70-80% confluence. After being changed to fresh medium for 1h, cells were treated with 10 mg/mL puromycin for 15 min. Elongation of the nascent polypeptide can be terminated by puromycin which incorporates into the synthesized polypeptide by mimicking the aminoacyl-end of an aa-tRNA. After being washed twice with cold PBS, cells were lysed with SDS-PAGE sample buffer. Western blot was used to detect the expression of puromycin by using an anti-puromycin antibody (ABclonal, A21205).

### RIP-qPCR

The RNA immunoprecipitation experiment was conducted according to manufacturer’s instructions with minor modifications. Briefly, 1×10^7^ HeLa cells in a 150-mm dish were subjected to RIP-qPCR. Cell pellets were lysed in RIP buffer (25 mM Tris, pH 7.4, 150 mM KCl, 2 mM EDTA, 0.5% NP40, 0.5 mM DTT, 1 X protease inhibitor cocktail). The mRNAs were then pulled down by antibody following the immunoprecipitation protocol. The co-precipitated RNA was recovered from the beads using a RNeasy mini kit (Qiagen Inc., Valencia, CA, USA). Both the input and co-immunoprecipitated RNA were recovered using an RNA Clean & Concentrator-5 kit (R1013; Zymo Research, Irvine, CA, USA) and subjected to RT-qPCR analyses. Primer sequences are provided in Supplementary Table 1.

### Ribo-seq

Six 15-cm plates of Hela cells at 80–90% confluency were used per group. Cells were treated with 100 μg/ml Cycloheximide for 8 minutes at 37°C, followed by lysis on ice for 10 minutes in 500 μl of lysis buffer (20 mM Tris-HCl pH 7.5, 150 mM NaCl, 5 mM MgCl₂, 1 mM DTT, 100 μg/ml Cycloheximide, and 1% Triton X-100). Lysates were cleared by centrifugation at 12000 g for 10 minutes at 4°C, and the absorbance at 260 nm was measured. Ribosomes from each group were normalized based on equal A₂₆₀ values and retained for RNA sequencing. RNase I (15 U per 1 OD of lysate) was added, and samples were incubated at 22°C for 40 minutes with gentle mixing. Ribosome-protected fragments (RPFs) were isolated using MicroSpin S-400 columns (Cytiva) and purified with the RNA Clean & Concentrator-25 kit (Zymo). RPFs were then separated on a 15% TBE-urea polyacrylamide gel, and RNA fragments between 17 and 34 nucleotides were excised and recovered using the ZR small-RNA™ PAGE Recovery Kit (Zymo Research). Purified RNA was treated with polynucleotide kinase (PNK), and libraries were prepared using the VAHTS™ Small RNA Library Prep Kit for Illumina®.

### Polysome profiling

Cells were pretreated with 100 μg/ml Cycloheximide (HY-12320; MedChemExpress) at 37°C for 8 minutes, washed with ice-cold PBS containing 100 μg/ml cycloheximide, and lysed on ice for 10 minutes in polysome lysis buffer (20 mM Tris-HCl pH 7.5, 150 mM NaCl, 5 mM MgCl₂, 1% Triton X-100, 100 μg/ml Cycloheximide, and 1 mM DTT). The samples were centrifuged at 13000 g for 15 minutes at 4°C, and the supernatants were collected for absorbance measurement at 260 nm. A 10-50% sucrose gradient was prepared using a Gradient Master (BioComp Instruments, Fredericton, Canada). Equal amounts of lysate, normalized based on A₂₆₀ values, were layered onto a 12mL sucrose gradient and subjected to ultracentrifugation at 35000 rpm for 3 hours at 4°C using an SW41Ti rotor (Beckman Coulter, Brea, USA). Following centrifugation, fractions were collected using a BioComp Piston Gradient Fractionator, and absorbance at 260 nm was recorded. The collected fractions were subsequently analyzed by western blot and RT-qPCR.

### Cell synchronization and cell cycle analysis

HeLa cells were synchronized at the G1/S-phase boundary using a double thymidine block protocol. HeLa WT and ZCCHC4 knockout cells were treated with thymidine (2 mM) for 16 hours. Following the first block, the thymidine-containing medium was removed and cells were washed three times with 15 mL of PBS to release the S-phase block. Cells were then cultured in fresh growth medium for 8 hours. Subsequently, the medium was replaced with growth medium containing an additional thymidine (final concentration 2 mM) for a second block lasting 16 hours. Cells were harvested at appropriate time points for cell cycle analysis. Cells were fixed in 70% ice-cold ethanol overnight and stained with FxCycle™ Violet (Thermo Fisher Scientific), which binds to double-stranded DNA, for 16 hours at 4°C. Stained cells were analyzed using a BD LSRFortessa™ flow cytometer (BD Biosciences) equipped with a 405 nm laser. For each sample, 1000 events were collected. FlowJo software (version 10.1, FlowJo LLC) was used to analyze data and determine cell cycle distribution using Watson’s pragmatic model^35^.

### Annexin V-PI staining

Apoptosis was assessed using an Annexin V-FITC apoptosis detection kit (A211; Vazyme) according to the manufacturer’s instructions. ZCCHC4 knockout HeLa cells were harvested, washed twice with cold PBS, and resuspended in Annexin V binding buffer. Subsequently, 5 µL of FITC-conjugated AnnexinV was added, and samples were incubated in the dark at room temperature for 15 minutes. For double staining, 5 µL of propidium iodide (PI, 5 mg/mL) was added 5 minutes before analysis. Apoptotic cells were quantified by flow cytometry using a FACS Aria cell sorter (BD Biosciences), with FITC fluorescence detected in the FL1 channel and PI fluorescence in the FL3 channel. A total of 10000 events per sample were recorded, and data were analyzed using FlowJo software.

### EdU assays

DNA synthesis was assessed using the Click-iT™ EdU Alexa Fluor™ 488 Imaging Kit (Thermo Fisher Scientific) following the manufacturer’s instructions. Briefly, cells were seeded in 6-well plates and cultured until they reached appropriate confluency. Cells were then incubated with 10 μM EdU for 2 hours at 37°C to label newly synthesized DNA. After EdU incorporation, cells were fixed with 4% paraformaldehyde for 15 minutes at room temperature and permeabilized using 0.5% Triton X-100 in PBS for 20 minutes. EdU detection was performed using the Click-iT reaction cocktail for 30 minutes in the dark. Nuclei were counterstained with DAPI (1 μg/mL) for 10 minutes. Images were acquired using a fluorescence microscope, and at least three random fields were captured per sample. The percentage of EdU-positive cells was calculated using ImageJ software (NIH), representing the fraction of cells undergoing DNA replication.

### Animals

To assess the tumor-suppressive role of ZCCHC4, a xenograft assay was conducted. A total of 1×10^7^ cells were suspended in an equal volume of Matrigel (Corning) and subcutaneously injected into the right flank of male nude mice (6-8 weeks old). Subcutaneous tumor formation was measured on day 9 and then every other day until the maximal tumor size reached 1600 mm^3^. The tumor volume is calculated as the product of length × width^2^. At endpoint, the tumors were collected to measure tumor weight. All mice were bred under specific-pathogen-free (SPF) conditions at the Animal Experiment Center of SUSTech. During this study, animal experiments were conducted in the Experimental Animal Center of Southern University of Science and Technology (SUSTech), research protocols were approved by the Institutional Animal Care and Use Committees (IACUC) of SUSTech.

### RNA-seq and ribo-seq analysis

Adapter sequences were removed from both Ribo-seq and RNA-seq reads using TrimGalore (v0.6.7). For Ribo-seq data, reads shorter than 20 nucleotides or longer than 35 nucleotides were discarded to ensure high-quality ribosome-protected fragment selection. The trimmed reads were then aligned to rRNA and tRNA sequences using Bowtie2 (v2.2.5)^53^. Reads that aligned to rRNA or tRNA were excluded, while the remaining unaligned reads were retained for subsequent analysis. Processed reads were aligned to the reference genome (gencode GRCh38) by the STAR (v2.7.10a)^54^, allowing for at most two mismatches. For Ribo-seq data, quality control and translational profiles was analyzed by RiboseQC. Transcriptome assembly and quantification were performed using StringTie (v2.1.7)^55^. Both read counts of Ribo-seq and RNA-seq were converted to TPM (transcripts per million). Genes with sufficient expression level (Ribo TPM > 1 and RNA TPM > 5) were subjected to further analysis. Translation efficiency (TE) and the differentially-TE genes (DTEGS) were calculated by RiboDiff (v0.2.2)^56^. GO term analysis were performed by Metascape (https://metascape.org/gp/index.html). GraphPad Prismsoftware and R was used for data presentation.

### ATAC-seq analysis

Raw paired-end reads were processed using Trim Galore for quality control and adapter trimming, during which low-quality bases and adapter contamination were removed. Then, reads were aligned to the reference genome using Bowtie2 (v2.2.5), and sorted BAM files were generated. Duplicate reads were marked using Picard, and low-quality, duplicated, and multimapped reads were filtered out using Samtools. ATAC shift correction was performed with alignmentSieve, and peaks were called using MACS2. The resulting BAM files were converted to bigWig format for signal visualization. Signal enrichment around transcription start sites (TSS) was visualized using computeMatrix and plotHeatmap/plotProfile. Finally, overlapping peaks between replicates were identified using Bedtools to generate a consensus peak set, and read coverage over these regions was quantified for downstream differential analysis.

### Statistical analysis

No statistical methods were used to predetermine sample size. Blinding and randomization were not applied, and no samples or animals were excluded from the analyses. All statistical analyses were conducted using GraphPad Prism 9 (GraphPad Software). For comparisons between two groups, two-tailed unpaired Student’s t-tests were performed. One-way analysis of variance (ANOVA) followed by Tukey’s multiple comparisons test was used for analyzing differences among more than two groups. Two-way ANOVA was employed for cell proliferation assays and tumor growth curve analyses. Statistical significance was considered at \**P*<0.05, \*\**P*<0.01, \*\*\**P*<0.001, \*\*\*\**P*<0.0001 or ns (not significant). The number of biological replicates is indicated in the figure legends.

## Acknowledgements

We would like to thank all members in Hao Chen’s Lab for their help and advice in experimental design. The authors would also like to acknowledge the technical support from Hua Li and Lin Lin at SUSTech CRF. This work was supported by Center for Computational Science and Engineering at Southern University of Science and Technology.

## Author Contributions

H.H.M., H.C. and Y.C.Z. designed and conceived the experiments. R.Q.W. and X.Y.S. performed most of the experiments. Y.Y.Z. helped to prepare figures. Y.L.C., C.C. and

R.F. assisted in polysome profiling. Y.C.W., X.Y.S. and H.C. analyzed the data.

R.Q.W. and H.C. wrote the manuscript. All authors have read and approved the final manuscript.

## Data and materials availability

**Extended Data Fig. 1:**
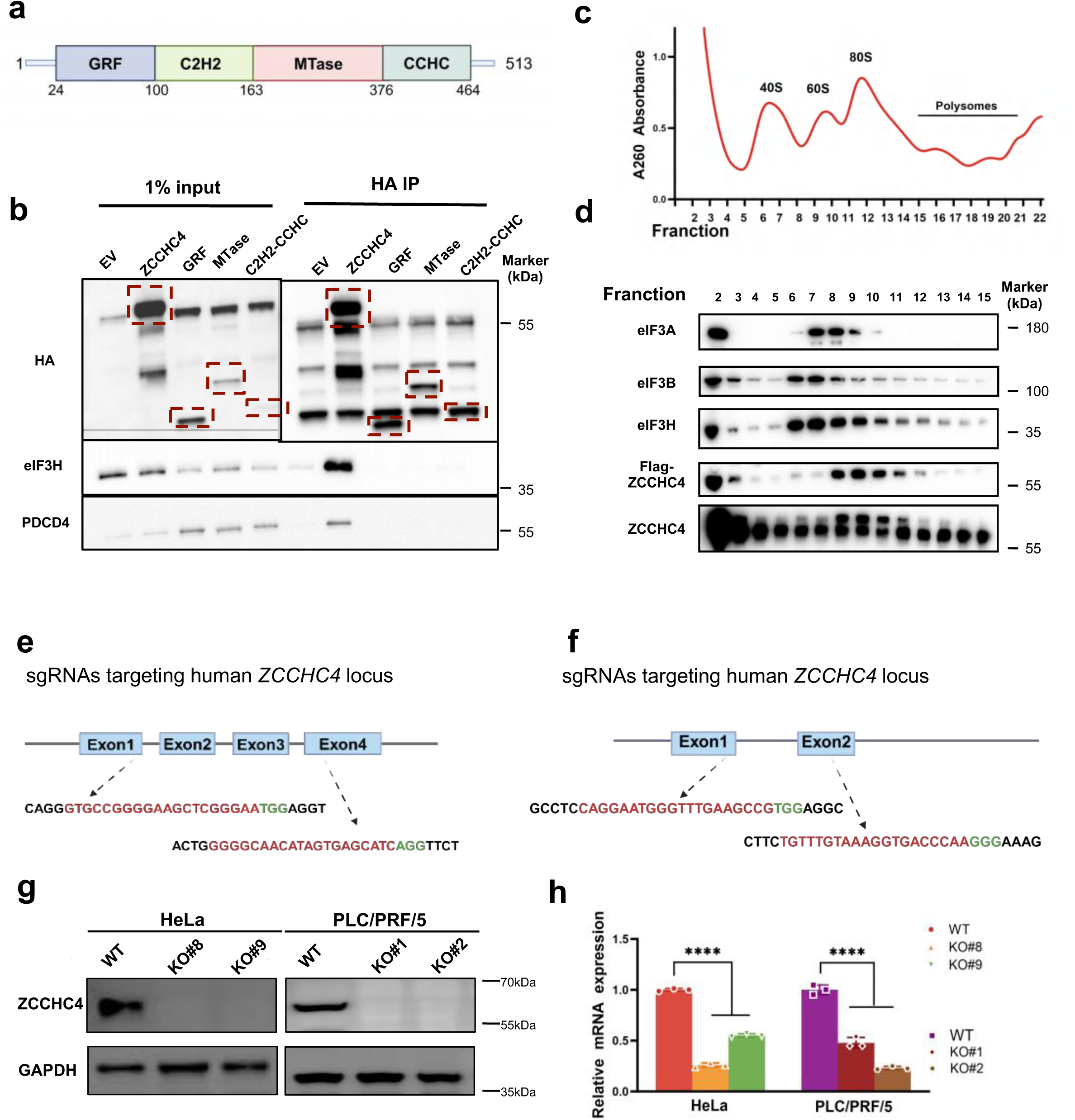
ZCCHC4 interacts with the eIF3 complex. **a,** Experimental strategy for constructing ZCCHC4 truncation mutants. **b,** Plasmid-expressing truncations of the GRF domain, MTase domain, and C2H2-CCHC zinc finger domain, were transfected into HEK 293T cells. HA beads were used to pull the complex containing ZCCHC4 and its interaction protein by immunoprecipitation in the cell lysate of HEK 293T cells. **c, d,** Polysome profiling of ZCCHC4 expression in HeLa cells (**c**). Western blot of eIF3A, eIF3B, eIF3H, ZCCHC4-Flag and ZCCHC4 demonstrating the components of the fraction after polysome profiling (**d**). **e, f,** Schematic representation showing the knockout strategy. Schematic diagram of sgRNA targeting human *ZCCHC4* in HeLa (**e**) and PLC/PRF/5 (**f**)cells. **g,** HeLa cells of WT and ZCCHC4 KO were established and confirmed by western blot. GAPDH was used as a loading control. **h,** RT-qPCR assays showed the mRNA expression of ZCCHC4 in HeLa and PLC/PRF/5 cells. **Note: h,** One-way ANOVA with Tukey’s test. Data were represented as mean ± SD (n=3). \*\*\*\**P* < 0.0001.

**Extended Data Fig. 2:**
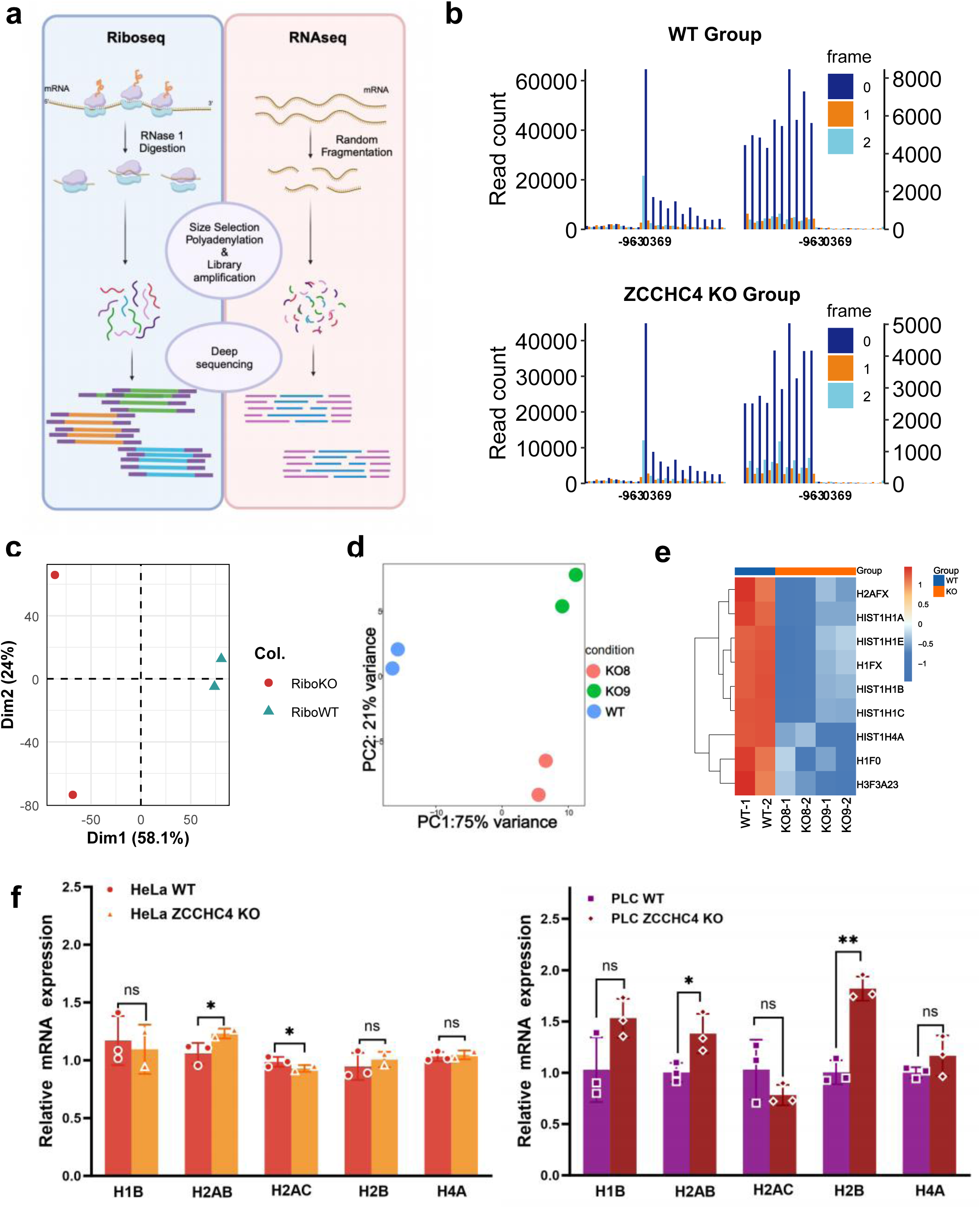
Multi-omics characterization of histone regulation by ZCCHC4. **a,** Schematic diagram of the Ribo-seq and RNA-seq. **b,** Quality control of Ribo-seq data in WT (up) and ZCCHC4 KO (down) cells. **c,** PCA quality control of Ribo data in WT and ZCCHC4 KO cells. **d,** PCA quality control of TMT data in WT and ZCCHC4 KO cells. **e,** Heatmap showing the expression profiles of histone-related genes in HeLa WT and ZCCHC4 KO cells. **f,** Analysis of histone mRNA transcription levels by RT-qPCR after ZCCHC4 knockout in HeLa (left) and PLC/PRF/5 (right) cells **Note: f,** Two-tailed unpaired Student’s t tests. Data were presented as mean ± SD (n=3). \**P*<0.05, \***\****P*<0.01, ns: not significant.

**Extended Data Fig. 3:**
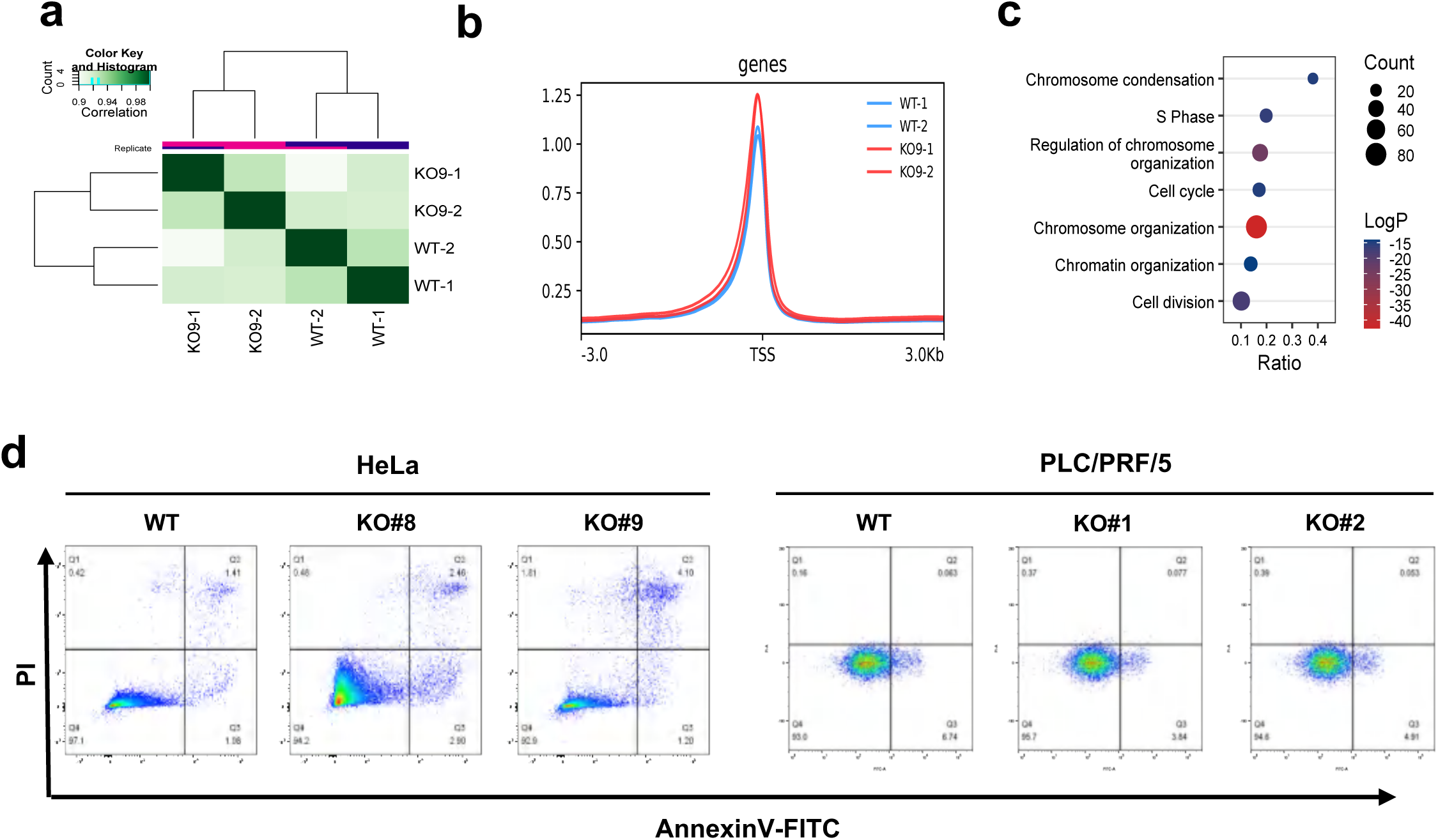
Effects of ZCCHC4 knockout on chromatin accessibility and apoptosis. **a,** Heatmap showing the Pearson correlations of two ATAC-seq replicates of the two sample groups. **b,** The average enrichment of ATAC-Seq peaks at annotated promoters (TSS ± 3kb) in HeLa WT and ZCCHC4 KO cells. **c,** GO analysis of TMT data in WT and ZCCHC4 KO cells. **d,** Flow cytometry of Annexin V and PI stain showed the apoptosis of HeLa and PLC/PRF/5 cells after ZCCHC4 knockout. FITC, Fluorescein 5-isothiocyanate; PI, Propidium Iodide.

**Extended Data Fig. 5:**
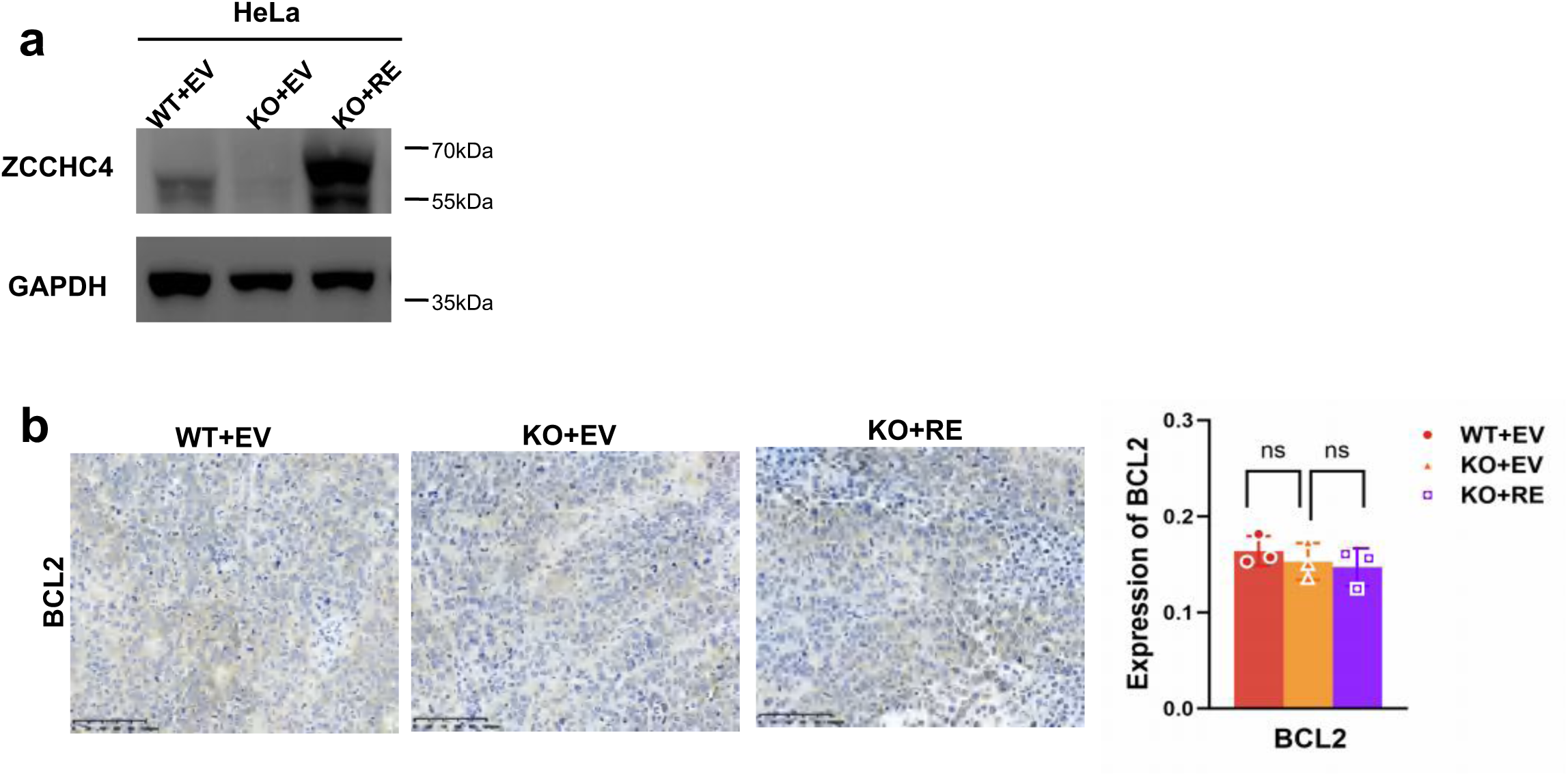
Deletion of ZCCHC4 impairs tumor progression. **a,** HeLa cells of WT, ZCCHC4 KO, and KO rescued with ZCCHC4 were established and confirmed by western blot. GAPDH was used as a loading control. **b,** Immunohistochemistry expression of BCL2 from WT, ZCCHC4 KO and KO rescued tumor. **Note: b,** Two-tailed unpaired Student’s t tests. Data were represented as mean ± SD(n=3). ns: not significant.

**Extended Data Fig. 6:**
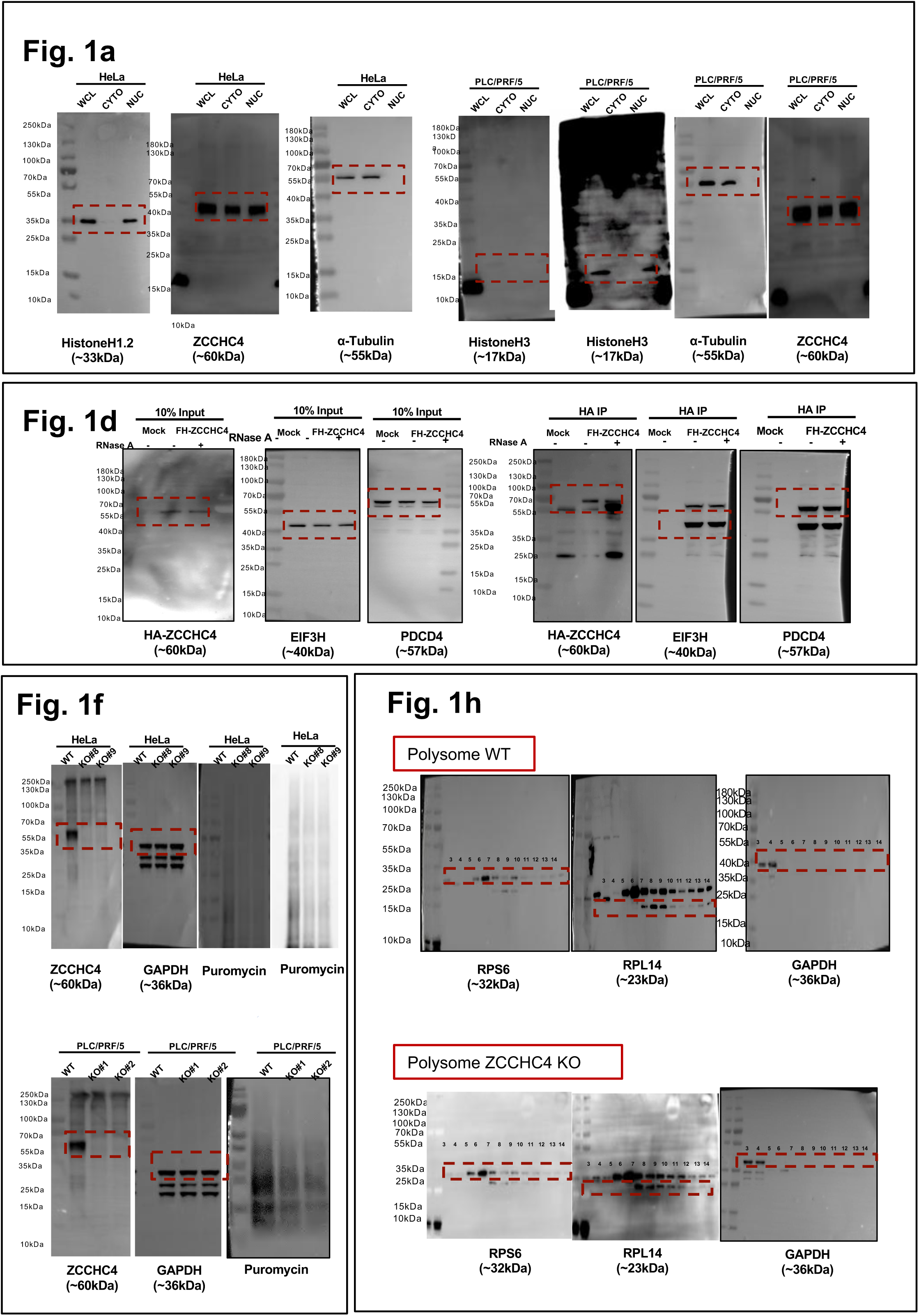
Original western blots.

**Extended Data Fig. 7:**
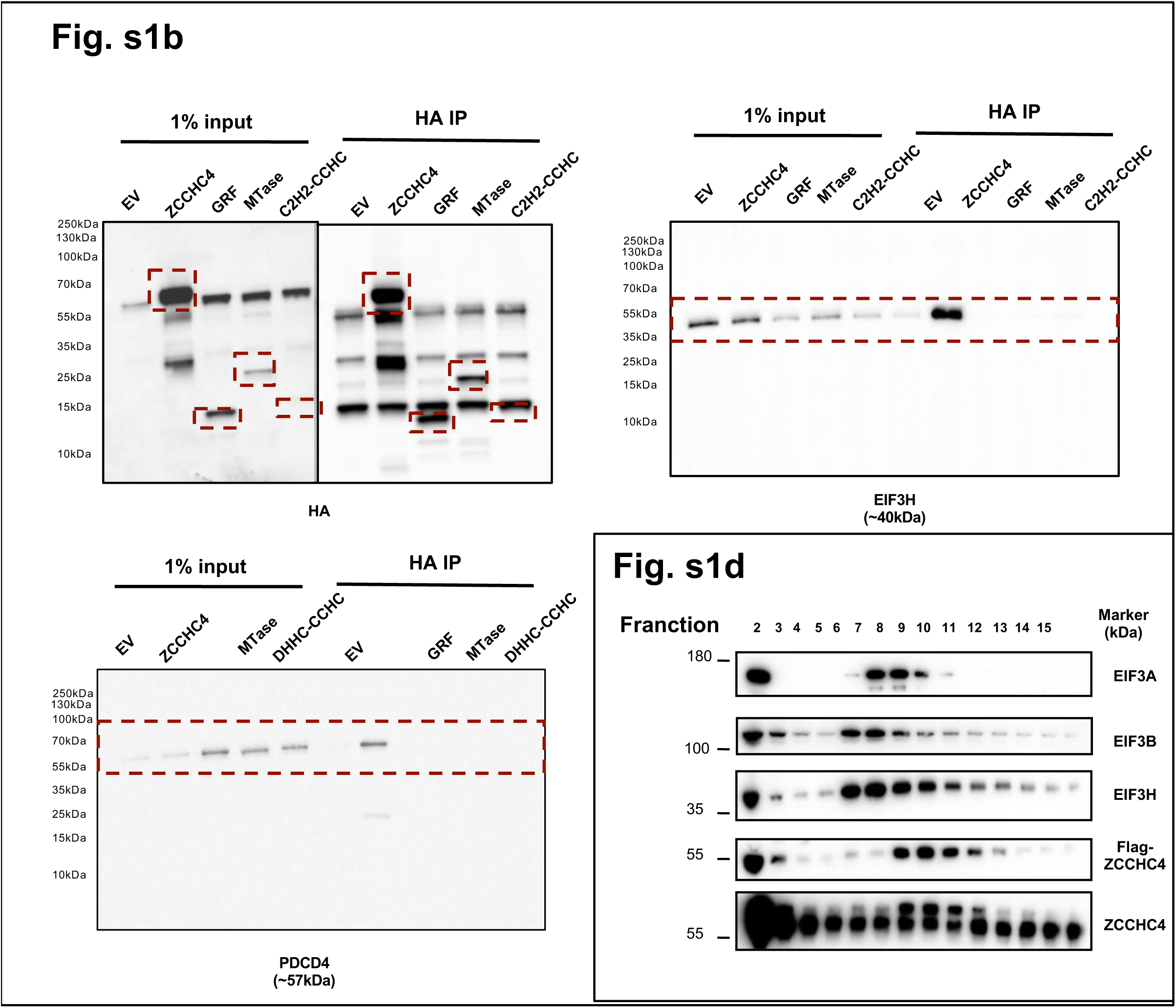
Original western blots.

**Extended Data Fig. 8:**
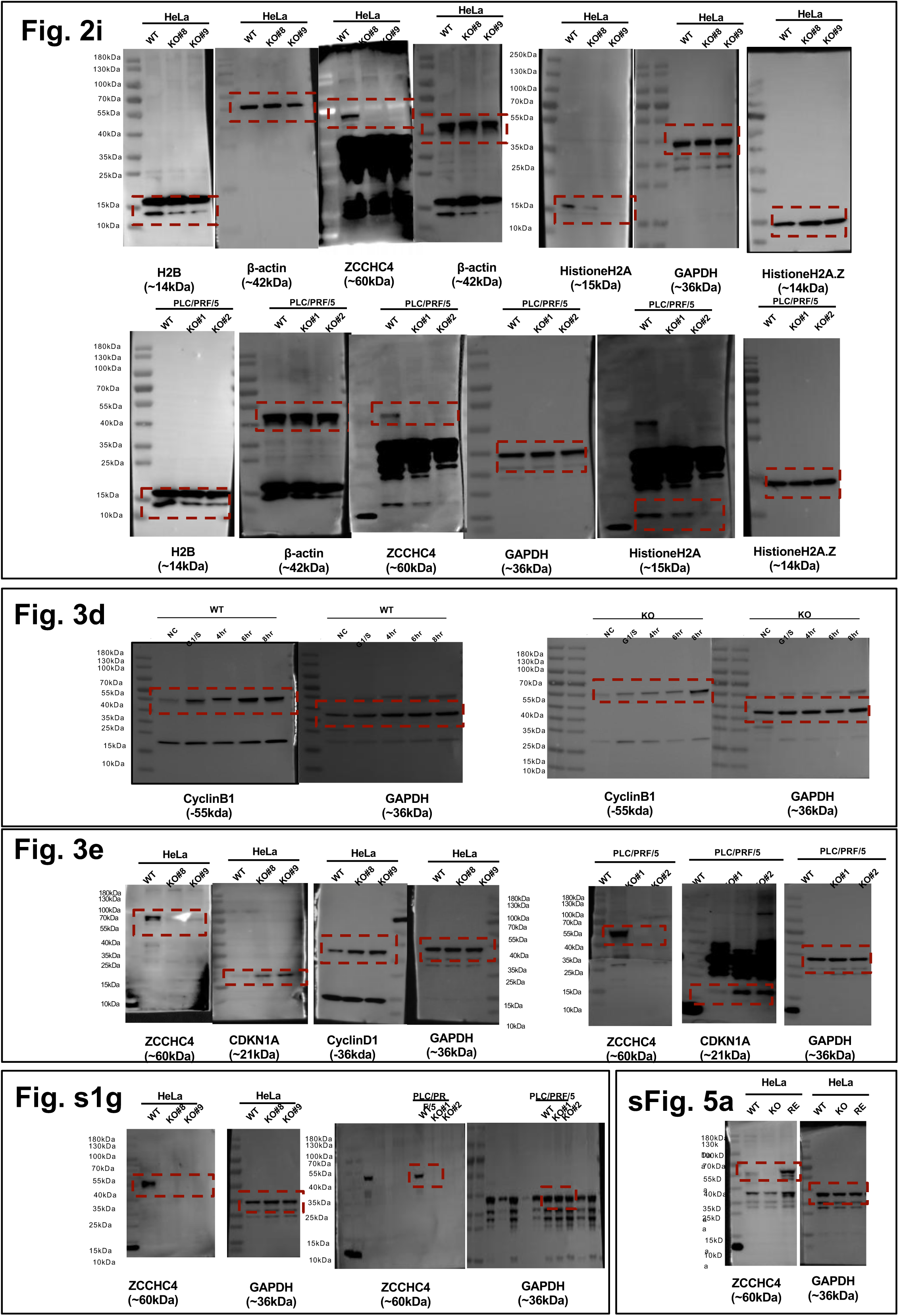
Original western blots.

**Supplementary Table S1.**
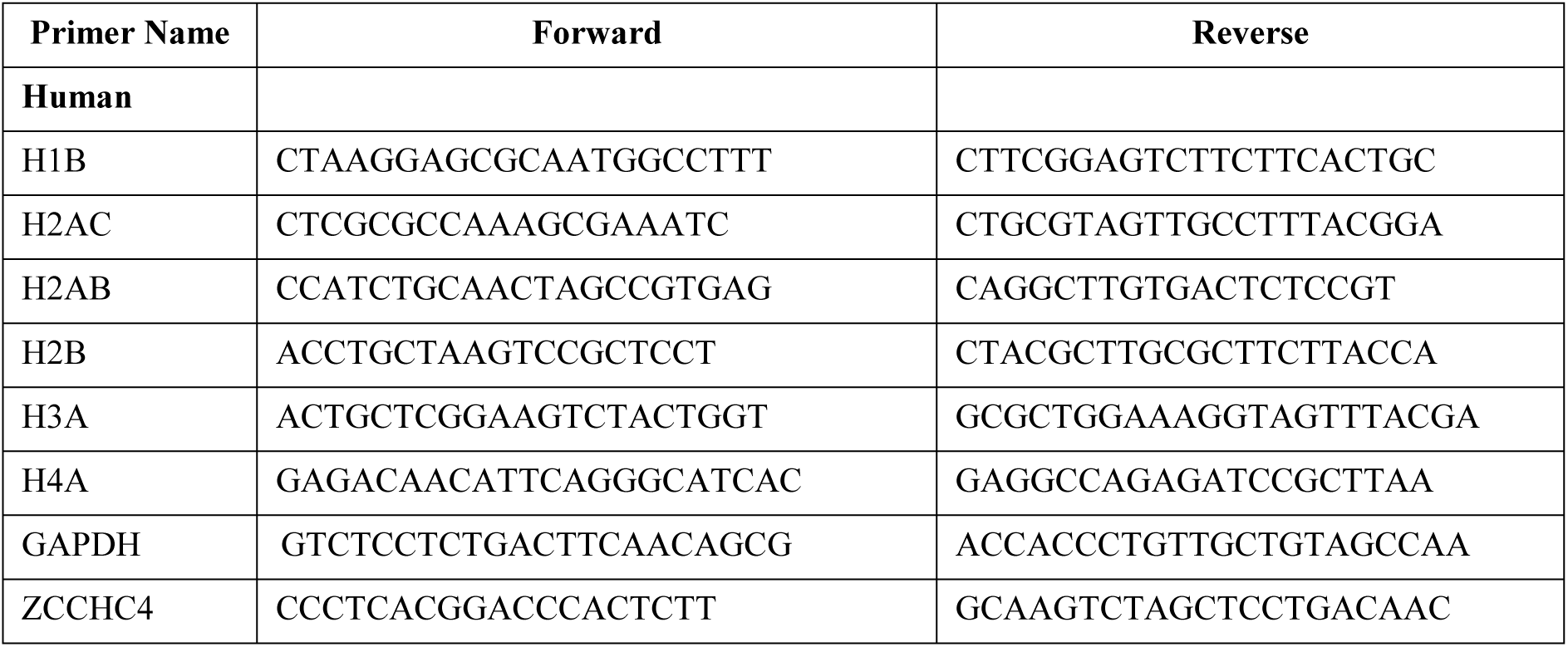
List of qPCR primers used in this study.

